# A High-Fidelity and Ancestrally Inclusive Patient-Derived Organoid Platform Resolves Cancer Cell Plasticity in Uterine Carcinosarcoma

**DOI:** 10.64898/2026.04.21.720027

**Authors:** Santhilal Subhash, Marie-Thérèse Bammert, Brian Yueh, Kadir A. Ozler, Timothy Chu, Melissa Kramer, Pascal Belleau, Astrid Deschênes, Onur Eskiocak, Vyom Shah, Charlie Chung, Ali Oku, Mali Barbi, Aybuke Alici, Megan Gorman, Arielle Katcher, Aaron Nizam, Ariel Kredentser, Divya Bhana, Jonathan Werner, Alexander M. Truskinovsky, Nicolas Robine, Marina Frimer, Gary L. Goldberg, Semir Beyaz

## Abstract

Uterine carcinosarcoma (UCS) is a rare but extremely lethal endometrial cancer that metastasizes early and resists current treatment modalities. It is biphasic, built from malignant epithelial and mesenchymal cells. Genomic studies indicate that these tumors are clonal, and that the mesenchymal cells arise from the epithelial cells through cancer cell plasticity. This biology has been hard to study, because faithful patient-derived models are scarce. The gap is widened by inequity. Women of African ancestry carry the greatest burden of UCS, yet are underrepresented in existing models. To address this, we established patient-derived organoids (PDOs) from an ancestrally inclusive UCS cohort, alongside matched normal endometrial PDOs. The organoids reproduced the biphasic histology of the original tumors. Across four sequencing platforms, they retained the tumor mutation and copy-number landscape, remained stable across passages, and expanded for up to 28 months. At single-cell resolution, UCS PDOs captured both malignant compartments and traced continuous transcriptional trajectories along the epithelial-to-mesenchymal axis, capturing patient-specific cancer cell plasticity. The models also nominated candidate vulnerabilities in proof-of-concept therapeutic testing. UCS PDOs were enriched for CREB-family transcriptional programs, and CREB inhibition reduced their viability. Combined FGFR and YAP inhibition outperformed either agent alone. Together, this work delivers a histologically, genomically, and transcriptionally faithful, ancestrally inclusive, and lineage-resolved UCS organoid platform for studying cancer cell plasticity and its vulnerabilities in an aggressive and inequitably burdened cancer.

## Introduction

Uterine carcinosarcoma (UCS) is an aggressive subtype of endometrial cancer defined by its biphasic architecture, comprising malignant epithelial and sarcoma-like components^1–3^. Although UCS represents a minority of endometrial cancers, it accounts for a disproportionate number of disease-related deaths due to rapid progression, early dissemination, and high recurrence rate. Standard-of-care treatment consists of surgery followed by platinum- and taxane-based chemotherapy^1,4–6^. However, therapeutic responses are frequently transient and relapse is common. Clinical management is largely derived from other high-grade endometrial or ovarian cancers, yet survival outcomes remain poor and there is no consensus on effective UCS-specific regimens, highlighting a significant unmet medical need^7–9^.

Despite its clinical aggressiveness, progress in understanding UCS biology has been limited by the lack of physiologically relevant models that capture the biphasic lineage composition and dynamic cell-state transitions. This gap is further widened by the underrepresentation of diverse patient populations in current preclinical and translational studies^10,11^, despite known disparities in UCS incidence, severity, and outcomes across ancestry, age, and metabolic context^12,13,14^. As a result, current models fail to reflect the full biological and clinical heterogeneity of UCS, restricting mechanistic insight and the identification of actionable therapeutic vulnerabilities. To address these challenges, there is a need for model systems that not only preserve the genomic and transcriptional architecture of primary tumors but also recapitulate their cellular heterogeneity, lineage plasticity, and treatment responses. In particular, models that demonstrate concordance with primary tumors at the level of mutational landscape, copy-number alterations, and cell-state-specific transcriptional programs are essential to enable functional interrogation and therapeutic testing.

Conventional two-dimensional cell lines fail to recapitulate the biphasic organization, intercellular signaling networks, and lineage heterogeneity that define primary tumors, while prolonged *in vitro* cultivation often selects for dominant clones, erasing patient-specific plastic states^15,16^. *In vivo* models, while informative, are constrained by interspecies anatomical and reproductive cyclic differences between rodents and humans, which limit full physiological translatability and fail to adequately capture the complexity and diversity of human endometrial tumors^17,18^. Patient-derived organoids (PDOs) provide a powerful framework to overcome these barriers by preserving three-dimensional architecture, tumor-intrinsic heterogeneity, and patient-specific molecular features while enabling functional interrogation of therapeutic responses in a controlled yet physiologically relevant context^19^. However, robust and inclusive organoid models of UCS that integrate genomic fidelity, lineage-resolved transcriptomics, and systematic chemotherapy modeling have not been comprehensively established^20–22^.

Recent efforts to develop patient-derived UCS models have underscored both the need for improved preclinical systems and the opportunity to address this gap. Patient-derived xenografts recapitulate some features of UCS but often engraft inefficiently and may fail to maintain the biphasic epithelial and mesenchymal architecture of the disease, particularly across serial passages^23^. More recently, UCS organoids have been used to test specific agents, including TROP2-directed antibody-drug conjugates^24^ and, in xenograft and cell-line models, FGFR1-directed inhibitors^25^, and organoids from a single patient have been used to study spatial heterogeneity^26^. These studies establish feasibility but have largely been built on ancestrally narrow cohorts and have not resolved, at single-cell resolution, the epithelial-to-mesenchymal (E/M) plasticity that defines UCS, nor benchmarked organoid fidelity across multiple orthogonal genomic platforms. A model system that is genomically faithful, ancestrally inclusive, and lineage-resolved is therefore still lacking.

Here, we establish an ancestrally inclusive cohort of UCS and matched normal PDOs that faithfully recapitulate the genetic, transcriptional, and cellular features of primary tumors. Using integrated whole-genome sequencing (WGS), whole exome sequencing (WES), targeted, and copy-number sequencing together with bulk and single-cell transcriptomics, and functional perturbation, we demonstrate high concordance between UCS PDOs and their tumors of origin, including preservation of biphasic epithelial and mesenchymal lineage states and inter-patient heterogeneity. We further show that these models resolve EMT trajectories, capture signaling dependencies and chemotherapeutic responses, enabling the identification of state-dependent vulnerabilities and providing a scalable platform for mechanistic studies. Together, this work closes a critical gap in UCS research by establishing a genomically and transcriptionally faithful, ancestrally inclusive PDO model system that enables the study of tumor plasticity, cell-state interactions, and therapeutic response, thereby advancing both biological understanding and translational opportunities in this lethal disease.

## Results

### Genomically defined and ancestrally inclusive UCS cohort enables PDO modeling

To establish a clinically relevant UCS model resource, we collected 25 UCS tumor specimens (23 primary tumors and 2 metastases) together with matched normal tissue when available and processed them for PDO derivation and molecular profiling (**Figure 1A**). Since UCS disproportionately affects women of African American (AFR) ancestry^27,28^ and prior The Cancer Genome Atlas (TCGA)-UCS studies included only 16 % (9/57) AFR patients^29^, our cohort was intentionally enriched for diversity, comprising 60 % (15/25) AFR women to address this health disparity. Patients were stratified by BMI, age, and self-reported ethnicity (**Figure 1B**). Whole genome sequencing (WGS) was performed on all 25 specimens to define somatic and germline alterations (**Figure 1C**). Genetic ancestry was determined from germline single nucleotide variants by projecting samples onto reference populations using ADMIXTURE^30^ inference. The predominant (max) ancestry for each specimen was defined as the ancestry component with the highest proportional contribution. Consistent with TCGA-UCS data^29^, *TP53* mutations were highly prevalent and detected in 21 of 25 tumors, underscoring the serous-like, copy-number high molecular architecture characteristic of UCS^31^. Additional mutations, including *ARID1A*, *PTEN*, *PIK3CA*, *FBXW7*, and *ASPSCR1*, varied across patients, reflecting substantial inter-tumoral heterogeneity. MANTIS^32^ analysis further identified one case (CS-014T) as microsatellite instability-high (MSI-H) with elevated tumor mutational burden (TMB), highlighting the presence of distinct molecular subsets within the cohort (**Figure 1C**).

**Figure 1:**
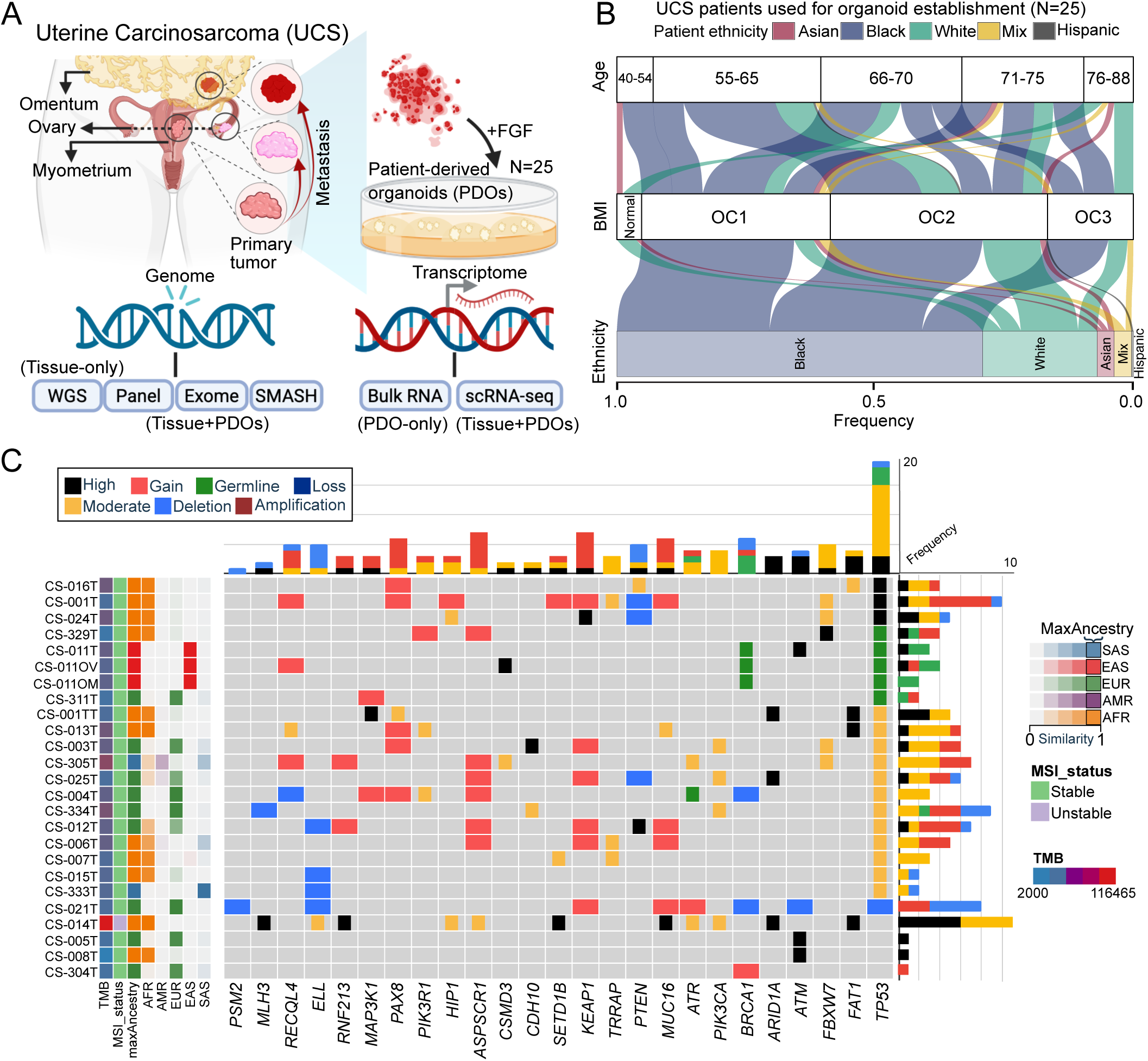
Genomic profiling defines an ancestrally enriched uterine carcinosarcoma cohort. (A) Schematic overview of sample collection and experimental workflow. Twenty-five tumor specimens from patients with uterine carcinosarcoma (UCS) were collected (n = 23 primary tumors; n = 2 metastases), with matched normal tissue when available, for the establishment of patient-derived organoids (PDOs). These UCS PDOs along with the primary tissues were sequenced using multiple genomic panels such as targeted Panel-seq (n = 36), Whole exome-seq (n = 8), and genome-wide copy-number profiling by SMASH-seq (n = 6). Whole genome sequencing (WGS) was only generated for primary tissues (n = 25). Single cell transcriptome was generated for both UCS PDOs and the primary tissues (n = 4). Bulk RNA-seq was only performed for PDOs (n = 17). (B) Alluvial plots show the overall distribution of UCS patients (n = 13) from which UCS PDOs are established in this study. This includes self-reported ethnicity (African American (AFR) n = 12, Asian (AS) n = 3, European/White (EUR) n = 6, mixed ethnicity (MIX) n = 3, and Hispanic n=1), body mass index (BMI), and age groups. The BMI groups are classified based on their BMI indices such as Normal >= 18.5 & < 25; Obese class 1 (OC1) >= 25 & < 30; Obese class 2 (OC2) >= 30 & < 40; Obese class 3 (OC3) >= 40. (C) Genomic landscape of the UCS cohort based on WGS. Heatmap displays recurrent somatic alterations across 25 UCS tissue specimens and tumor mutational burden (TMB). Microsatellite instability (MSI) status was assessed using MANTIS. Ancestry inference was performed using ADMIXTURE analysis and was concordant with self-reported ethnicity. AFR: African American; AS: Asian; EUR: European/White; EAS: East Asian; SAS: South Asian.

Collectively, this ancestrally enriched and genomically characterized UCS cohort provides a valuable and diverse resource that captures both canonical *TP53*-driven biology and patient-specific heterogeneity for organoid-based studies.

### FGFR genomic alterations inform FGF-dependent establishment and long-term expansion of UCS PDOs

We next asked whether recurrent growth factor signaling alterations in UCS could help guide culture conditions for robust PDO establishment. UCS PDOs were established and expanded according to previously published protocols^33–35^, followed by comprehensive genomic, transcriptomic, and histologic profiling to interrogate UCS pathophysiology and identify candidate molecular targets. Analysis of the TCGA-UCS cohort (n = 56) revealed recurrent alterations in growth factor receptor genes, with the most prominent events involving *FGFR3*, *FGFR1*, and *FGFR2*, along with less frequent alterations in *PDGFRA*, *PDGFRB*, and *IGF1R* (**Figure 2A**). Because these events include amplification, mutation, and deletion, our data suggest a pathway-level dysregulation rather than a single recurrent hotspot event.

**Figure 2:**
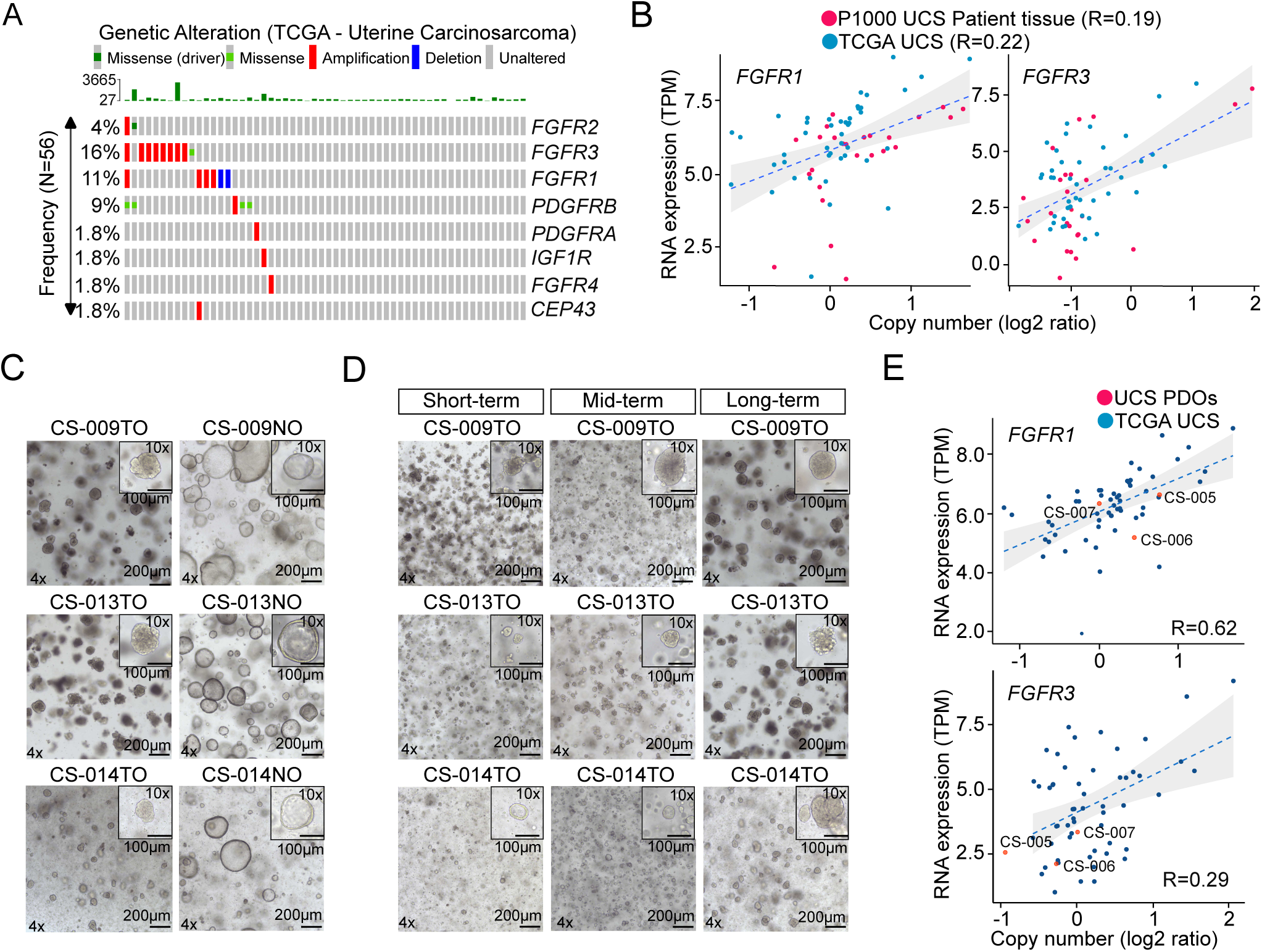
Recurrent FGFR alterations support FGF-dependent establishment of UCS PDOs. (A) Frequency and spectrum of recurrent genetic alterations in receptor tyrosine kinase and growth factor signaling genes in UCS from the TCGA cohort (n = 56). Alterations include non-synonymous somatic mutations and high-level copy-number gains or deletions. Thresholds to assign a significant alteration were set to p-value < 0.001 for deletions and p-value > 0.999 for amplifications. (B) Copy-number-expression correlation analysis for *FGFR1* and *FGFR3* in primary UCS tumors from our cohort (red; n = 25) compared with TCGA-UCS samples (blue; n = 56). Log2 copy-number ratios derived from whole-genome sequencing were plotted against transcript abundance (TPM) from bulk RNA sequencing. Correlation coefficients were calculated using Spearman method. (C) Representative brightfield images of matched normal endometrial and UCS PDOs for donors CS-009, CS-013, and CS-014. Scale bars: x4: 200 µm; x10: 100 µm. (D) Representative brightfield images of the long-term expansion of representative UCS PDO lines (CS-009, CS-013, and CS-014) maintained in culture for up to 28 months. Images show organoid morphology at indicated time points. Short-term:0.5-1.5 months; Mid-term: 5-8 months; Long-term: 18-28 months. Scale bars: x4: 200 µm; x10: 100 µm. (E) Copy-number-expression correlation for *FGFR1* and *FGFR3* in matched UCS PDOs (red, n = 3) compared with TCGA-UCS samples (blue; n = 56). Log2 copy-number ratios were derived from whole-exome sequencing and plotted against bulk RNA expression (TPM). Correlation was assessed using Spearman method.

To test whether genomic alteration translated into transcriptional output, we compared copy-number and RNA expression measurements for *FGFR1* and *FGFR3* in our cohort (n = 25, red) and in TCGA-UCS^29^ (n = 56, blue) (**Figure 2B**). In both, increased copy number was associated with higher transcript abundance, although the effect sizes were modest, which is expected for pathway components influenced by both dosage and cell-state context (**Figure 2B**).

Based on these observations, we hypothesized that FGF signaling may provide critical growth support for UCS cells *in vitro*. We therefore systematically tested individual FGF ligands and combinations during PDO establishment. Among tested conditions, FGF1, FGF4 and FGF9, ligands known to signal predominantly through FGFR1 and FGFR3^36^, robustly supported PDO formation either individually or in combination (**Figure S2A-B**). Incorporation of FGF1 and FGF4 into an optimized culture system resulted in successful establishment of UCS PDOs with an overall efficiency of approximately 85 % across donors (**Figure S2C**).

Brightfield imaging demonstrated stable three-dimensional architecture of UCS PDOs, which were morphologically distinct from matched normal endometrial PDOs derived from the same patients (**Figure 2C** and **Figure S2D**). Importantly, we were able to expand UCS PDO models long-term in culture for up to 24 months. (**Figure 2D** and **Figure S2E**). Quantification further showed an increase in PDO yield between early and later culture time points, indicating that PDO-forming cells remain proliferative and undergo expansion over time in culture (**Figure S2F**). A subset of these UCS PDOs were expanded for the Human Cancer Models Initiative for distribution with the scientific community^37^.

To evaluate whether tumor-intrinsic genomic features were preserved *in vitro*, we performed matched whole exome sequencing (WES) and bulk RNA-seq on three established UCS PDOs. We integrated UCS PDO transcriptomic profiles with their corresponding copy-number landscapes and conducted the same copy-number-expression correlation analysis as in primary tumors (**Figure 2E**). Consistent with patient tissue and TCGA-UCS samples, *FGFR1* and *FGFR3* copy-number gains in UCS PDOs were associated with increased transcript abundance (**Figure 2E**).

Collectively, these findings demonstrate that FGFR pathway is altered in UCS and incorporating FGFs support the robust establishment and long-term expansion of UCS PDOs.

### UCS PDOs recapitulate histopathologic and phenotypic features of primary tumors

To assess whether UCS PDOs preserve histopathological features of the parental tumors, we performed comparative hematoxylin and eosin (H&E) staining and immunohistochemical (IHC) analyses on matched primary tissues and PDOs. Primary UCS specimens displayed characteristic biphasic morphology, with malignant epithelial glandular structures intermingled with atypical mesenchymal component (**Figure 3A**). High-grade nuclear atypia and stromal expansion were evident across cases (**Figure 3A**, highlighted by asterisk and arrows).

**Figure 3:**
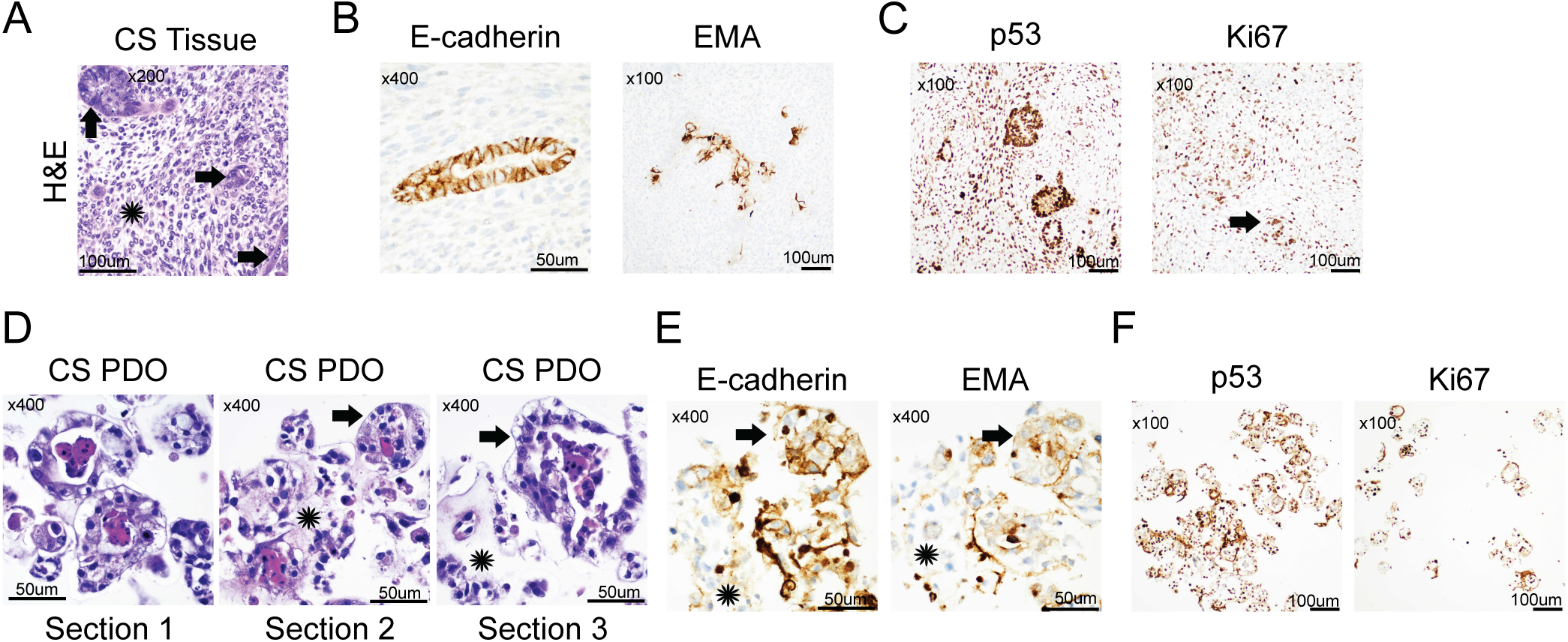
UCS PDOs retain the histologic and immunophenotypic features of primary tumors. (A) Representative images of hematoxilin & eosin (H&E) staining of primary UCS tumor tissue section showing malignant glands (arrows) surrounded by highly atypical stroma (asterisk), representing malignant mesenchymal differentiation of the tumor. Original magnification x400. Scale bar: 100 µm. (B) Representative immunostaining of UCS PDOs for E-cadherin and epithelial membrane antigen (EMA). The epithelial organoids are positive for E-cadherin, while the surrounding neoplastic stroma shows negative staining. Original magnification x400. Scale bar: 50 µm. Similar, EMA staining shows that epithelial elements are multifocally positive, while the malignant stroma is negative. Original magnification x100. Scale bar: 100 µm. (C) Representative immunostaining of UCS PDOs for p53 shows diffusely positive staining, including the neoplastic stroma in the uterine tumor. Scale bar: 100 µm. Representative immunostaining of UCS PDOs for Ki67 shows markedly increased proliferative index. Both the neoplastic glands (like the one marked by the arrow) and the surrounding neoplastic stroma are diffusely positive. Original magnification x100. Scale bar: 100 µm. (D) Representative H&E UCS PDO sections (S1, S2, and S3), consisting of the malignant epithelial elements of the tumor, show high-grade nuclear atypia and central necrosis (S1). Other areas show PDOs (arrow) with adjacent neoplastic cells not arranged in epithelial-like structures (star) (S2). In addition, an area suggestive of extracellular matrix formation is seen in (S3). Original magnification x100. Scale bar 100 µm. (E) Representative immunostaining of UCS PDOs for E-cadherin. The epithelial PDOs (arrow) show positive staining, while the adjacent disorganized neoplastic cells are negative (star). Scale bar: 50 µm. Representative immunostaining of the organoids for EMA shows that the epithelial elements (arrow) are multifocally positive, while the adjacent disorganized tumor cells are negative (star). Original magnification x400. Scale bar: 50 µm. (F) Representative immunostaining of UCS PDOs for p53 shows diffusely positive signals, including the neoplastic stroma. Original magnification: x100. Scale bar: 100µm. Representative immunostaining of Ki67 shows increased signal. Both the neoplastic glands (arrow) and the surrounding neoplastic stroma are diffusely positive. Original magnification x100. Scale bar: 100 µm.

Immunohistochemical staining of primary UCS tumors confirmed epithelial differentiation by E-cadherin (CDH1) positivity and epithelial membrane antigen (EMA) expression (**Figure 3B**), while diffuse p53 staining was observed, consistent with the high frequency of *TP53* alterations identified genomically (**Figure 3C**). A high Ki67 signal further indicated high proliferative activity within both epithelial and stromal compartments (**Figure 3C**).

Histological examination of UCS PDOs demonstrated preservation of malignant architectural features^38^, including high-grade nuclear atypia and regions of central necrosis (**Figure 3D**, **Figure S3A**-**D,** highlighted by asterisks and arrows). UCS PDOs displayed complex three-dimensional organization, and in some regions, extracellular matrix-like structures were observed, suggestive of stromal-like differentiation (**Figure 3D**, **Figure S3A**-**D**). These findings indicate that PDOs maintain structural features reminiscent of the biphasic nature of UCS.

Immunostaining of UCS PDO sections revealed cellular marker expression patterns comparable to the corresponding primary tumors. CDH1 confirmed epithelial components within PDOs, EMA marked epithelial differentiation, diffuse p53 expression reflected retention of *TP53*-altered tumor identity, and Ki67 staining demonstrated sustained proliferative capacity *in vitro* (**Figure 3E-F** and **Figure S3A-D**). Together, these data demonstrate that UCS PDOs preserve key histopathological characteristics of the parental tumors, supporting their suitability as clinically-relevant models.

### UCS PDOs preserve the genomic landscape and copy-number architecture of primary tumors

To investigate in genomic concordance between primary tumors and matched UCS PDOs, we performed multi-layered genomic profiling across sequencing platforms. Targeted panel sequencing of 49 recurrent cancer-associated genes was conducted on 25 tumor tissues (23 primary tumors and 2 metastases) and matched UCS PDO lines (**Figure 4A**, **Figure S4A**). Across samples, recurrent alterations were detected in canonical carcinosarcoma-associated genes, including *TP53*, *KRAS*, *PIK3CA*, *ARID1A*, and *TGFB2* (**Figure 4A**, **Figure S4A**), consistent with prior genomic studies^10,29,39,40^ and UCS primary tumor WGS (**Figure 1A**). Matched PDOs retained the vast majority of somatic mutations detected in their parental tumors (**Figure 4A**, **Figure S4A**). For example, a frameshift deletion in *ARID1A* was detected in both primary tumors and matched UCS PDOs (**Figure 4A**).

**Figure 4:**
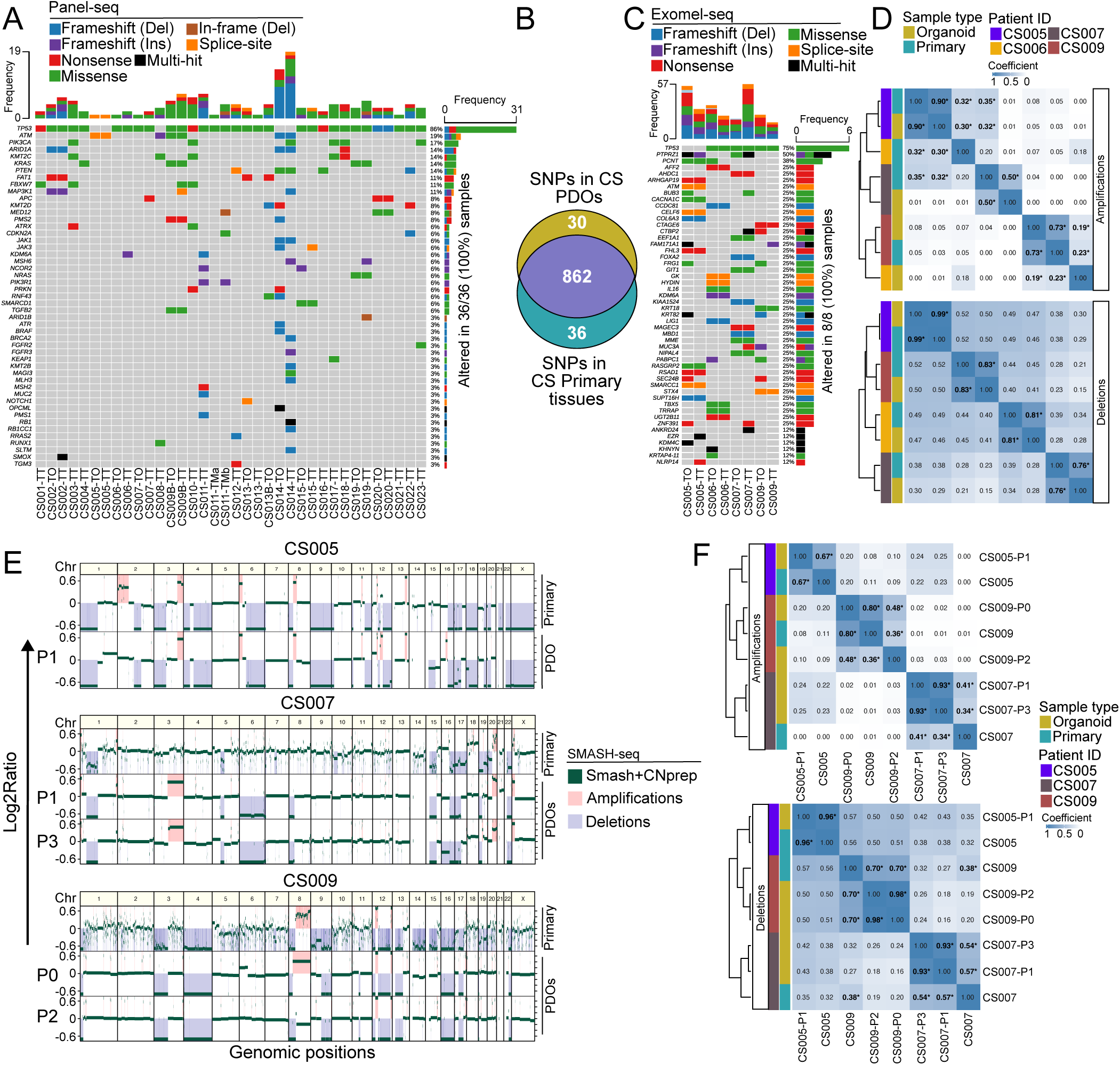
Preservation of mutational and copy-number landscapes in UCS PDOs. (A) Targeted panel-seq of 49 cancer-associated genes in UCS tumors (n = 25; 23 primary tumors, 2 metastases) and matched PDOs (n = 10 matched pairs). Oncoplot displays distribution of somatic mutations across samples. (B) Venn diagram illustrates the overlap of detected single nucleotide polymorphisms (SNPs) within the targeted regions between primary tumors and matched PDOs, comprising 862 shared variants, 36 primary-specific SNPs, and 30 PDO-specific SNPs. (C) Whole-exome sequencing (WES) oncoplot shows comparison of somatic mutational profiles between matched tumor-PDO pairs (n = 4 matched donors). (D) Heatmaps showing the Sørensen similarity indexes between CS tumors (TT) and PDOs (TO) exome profiles of chromosomal aberrations (Deletion and Amplification). In the heatmap value of 0 indicates no similarity, while 1 indicates exact similarity. * indicates the similarity index is significant with p-value < 0.05 calculated using simulations. (E) Genome-wide copy-number profiles generated by SMASH sequencing for primary tumors and matched PDOs across multiple passages. Log2 copy-number ratios are plotted across chromosomes. Regions with significant amplification and deletion events are respectively colored in pink and light blue. (F) Heatmaps showing correlation in amplification and deletion profiles between matched tumor-PDO pairs was performed using Sørensen correlation test. In the heatmap value of 0 indicates no similarity, while 1 indicates exact similarity. The * indicates the similarity index is significant with p-value < 0.05 calculated using simulations.

To quantify overlap between primary tumors and UCS PDOs, we compared single nucleotide polymorphisms (SNPs) identified across matched samples. Of the detected variants, 862 SNPs were shared between primary tissues and UCS PDOs, whereas 36 were exclusive to primary tumors and 30 were unique to UCS PDOs (**Figure 4B**).

To complement targeted genomic profiling, we performed whole-exome sequencing (WES) on three matched primary UCS tumor-PDO pairs (eight total lines including independent passages) to evaluate broader genomic concordance (**Figure 4C-D**, **Figure S4B**-**C**). Somatic mutation spectra were highly similar between matched pairs, and hierarchical clustering based on variant profiles grouped PDOs with their respective parental tumors. Importantly, analysis of copy-number alterations demonstrated concordance in both amplification and deletion events across matched samples, with high pairwise correlation coefficients (**Figure 4C-D**).

Given that UCS is characterized by widespread copy-number instability^29^, we next performed Short Multiply Aggregated Sequence Homologies (SMASH) sequencing to generate genome-wide copy-number profiles. SMASH analysis confirmed that UCS PDOs maintained large-scale chromosomal amplifications and deletions observed in the corresponding primary tumors (**Figure 4E**). Moreover, copy-number landscapes were stable across serial passages, indicating genomic stability during long-term *in vitro* culture. Correlation analyses of amplification and deletion profiles further demonstrated that UCS PDOs clustered with their matched primary tumors rather than with unrelated samples (**Figure 4F**).

Collectively, these multi-platform genomic analyses demonstrate that UCS PDOs faithfully preserve the mutational landscape and copy-number architecture of their parental tumors, supporting their use as a genomically stable model.

### Single-cell transcriptomics reveals cellular heterogeneity and epithelial-mesenchymal plasticity in UCS PDOs

To assess whether UCS PDOs preserve the cellular heterogeneity and lineage complexity of primary tumors, we performed single-cell RNA-seq on four independent UCS PDO lines (CS-007TO, CS-009TO, CS-013TO, and CS-014TO). UMAP embedding revealed distinct donor-specific clustering, reflecting inter-patient transcriptional heterogeneity (**Figure 5A**, **Figure S5A**-**B**). Within each UCS patient, cells segregated into carcinoma-like and sarcoma-like compartments, with further subdivision into epithelial basal, glandular, and luminal states, as well as endothelial-like, fibroblast-like, stromal-like, and muscle-like mesenchymal populations (**Figure 5A**). Additional stem-associated clusters, including mesenchymal stem cell (MSC)-like and cancer stem cell (CSC)-like populations, were identified (**Figure 5A**). The relative abundance of these cell states varied markedly across donors (**Figure 5A, lower panel**). Certain cell populations appeared selectively enriched in individual PDOs. For example, fibroblast-like cells were predominantly observed in CS-007TO, whereas CS-009TO displayed an expanded epithelial luminal compartment (**Figure 5A, lower panel**). Donor CS-014TO exhibited a distinct EMT-enriched subpopulation, while MSC-like states were present across donors, albeit with heterogeneous transcriptional signatures (**Figure 5A, lower panel**). These findings indicate that UCS PDOs capture both epithelial and mesenchymal lineages and reflect patient-specific cellular architecture.

**Figure 5:**
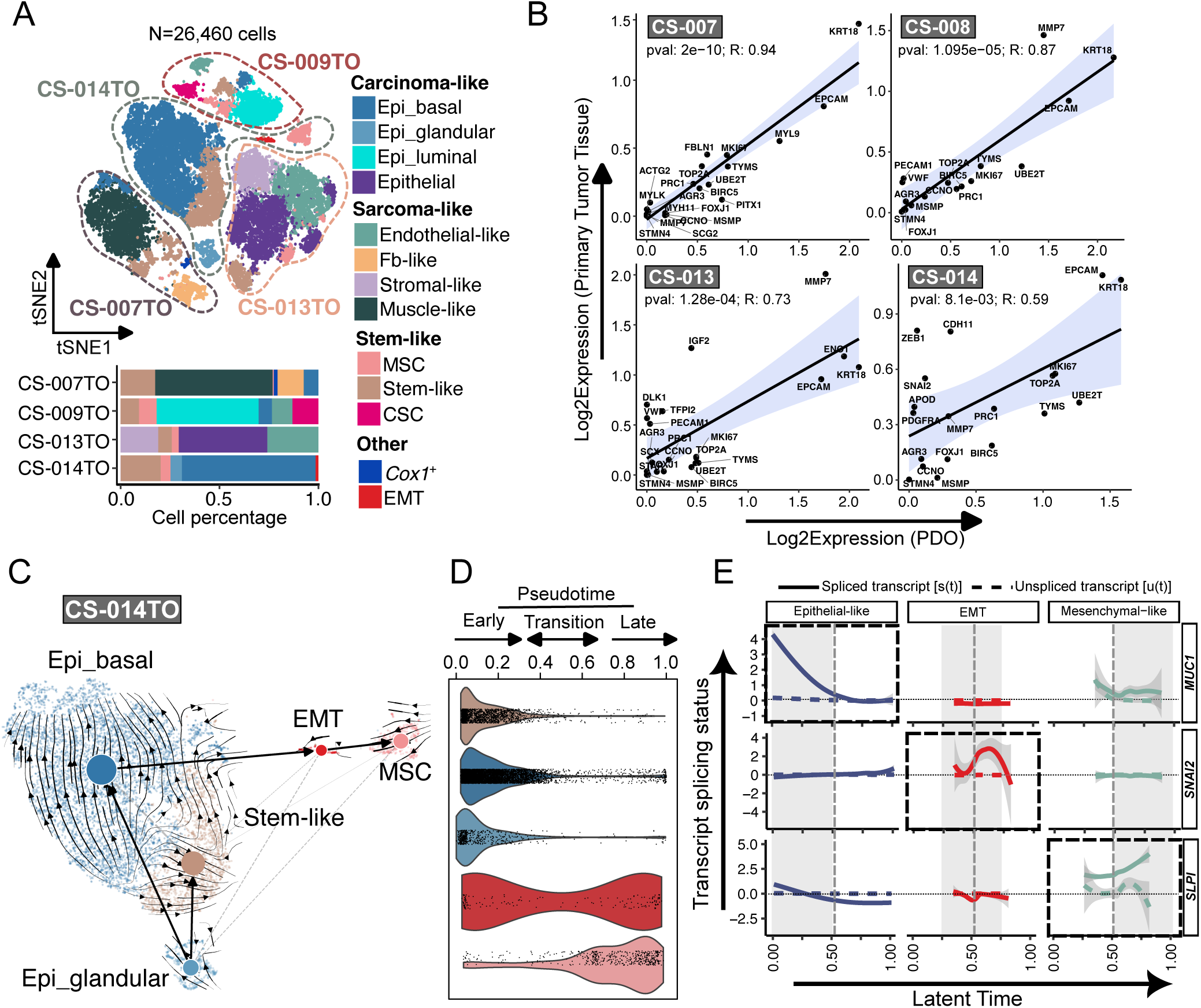
UCS PDOs preserve dynamic epithelial-mesenchymal transitions. (A) t-SNE representation of single-cell RNA sequencing (scRNA-seq) data (N= 51,784 cells) from four independent UCS PDO lines (CS-007TO, CS-009TO, CS-013TO, CS-014TO). Cells are colored by annotated cell subtype, including carcinoma-associated (epithelial basal, glandular, luminal, and epithelial populations), sarcoma-associated (endothelial-like, fibroblast-like (Fb-like), stromal-like, and muscle-like), stem-associated (mesenchymal stem cells (MSCs), stem-like, cancer stem cells (CSCs)), as well as other (*Cox1^+^*, EMT). Bar plot (below) shows proportional representation of each cell type per donor. (B) Correlation analysis of total gene expression signatures between matched primary tumors and PDOs. Each point represents a UCS donor. X-axis consists of log2 transformed expression in UCS PDOs and Y-axis consists of log2 transformed expression in primary UCS tumors. The correlation and p-values values are calculated using Pearson correlation coefficient. (C) RNA velocity analysis of representative PDO CS-014TO, showing directional transcriptional flow from epithelial basal and glandular populations toward EMT and stem- and MSC-like states. Velocity vectors were computed using scVelo based on spliced and unspliced transcript counts. (D) Pseudotime of transcripts from individual cells of PDO CS-014TO is inferred using scVelo, indicating early epithelial, transitional, and late mesenchymal states along a continuous trajectory. Pseudotime represents an unobserved, relative distance along a trajectory or differentiation time from one point to another (here, 0 to 1), inferred via transcriptomic similarity in individual cells. (E) Latent time derived from RNA velocity of PDO CS-014TO, illustrating dynamic progression across lineage states. Representative genes (*MUC1*, *SNAI2*, *SLPI*) are plotted across latent time to demonstrate sequential activation of epithelial, EMT, and mesenchymal programs.

To determine whether UCS PDOs maintain transcriptional similarity to primary tumors, we compared cell type-defining gene expression signatures between PDOs and matched tissues. Correlation analyses demonstrated concordance between UCS PDOs and primary tumor transcriptomes across donors (R = 0.59-0.94; **Figure 5B**), supporting preservation of tumor-intrinsic transcriptional programs.

We next analyzed transcriptional dynamics within UCS PDOs. Representative RNA velocity analysis in donor CS-014TO revealed directional flow from epithelial basal and glandular populations toward EMT and MSC-like states (**Figure 5C**). Pseudotime analysis further resolved cells along an early epithelial, transitional, and late mesenchymal continuum (**Figure 5D**). Latent time modeling confirmed a continuous trajectory, with gene-specific splicing kinetics indicating temporal activation of EMT-associated programs (**Figure 5E**). For example, canonical epithelial markers (e.g., *MUC1*) were enriched in early epithelial states, whereas EMT regulators (e.g., *SNAI2*) peaked during transition, and mesenchymal markers (e.g., *SLPI*) were upregulated at later stages (**Figure 5E**). Comparable trajectory architectures were observed across additional UCS PDOs (CS-007TO, CS-009TO, and CS-013TO) (**Figure S5C**). In each donor, RNA velocity streams indicated directional progression from epithelial or stem-associated states toward mesenchymal-like compartments, although the dominant originating populations varied between patients (**Figure S5C**).

Collectively, these analyses demonstrate that UCS PDOs recapitulate patient-specific cellular heterogeneity while preserving dynamic epithelial-mesenchymal state transitions.

### Integrated transcriptomic profiling identifies CREB-associated transcriptional programs in UCS PDOs

To define UCS PDO specific transcriptional programs compared to normal PDOs, we performed bulk RNA-seq on eight UCS PDO lines and nine normal or matched endometrial biopsy-derived PDOs. Principal component analysis demonstrated segregation between UCS and normal samples (**Figure S6A**), confirming distinct transcriptomic identities. ADMIXTURE^41^-based ancestry inference from bulk RNA-seq further validated that the majority of sequenced samples were of African ancestry (**Figure S6B**).

Differential expression analysis identified 2,158 significantly deregulated genes in UCS PDOs relative to normal PDO controls (FDR < 0.05), including 1,336 upregulated and 822 downregulated transcripts (**Figure 6A**). Among the most significantly upregulated genes were proliferation- and EMT-associated transcripts (e.g., *KIF5C*, *S100A9*, *SIX1*, *IGF2*), whereas downregulated genes included epithelial-lineage regulators and differentiation-associated transcripts (**Figure 6A**). Multiple long non-coding RNAs (lncRNAs) were significantly deregulated, indicating coordinated remodeling of both coding and non-coding transcriptional networks (**Figure 6A**). In addition, FGFR expression was altered among UCS compared to normal PDOs concordant with our observations on a genomic level (**Figure S6C**-**D**).

**Figure 6:**
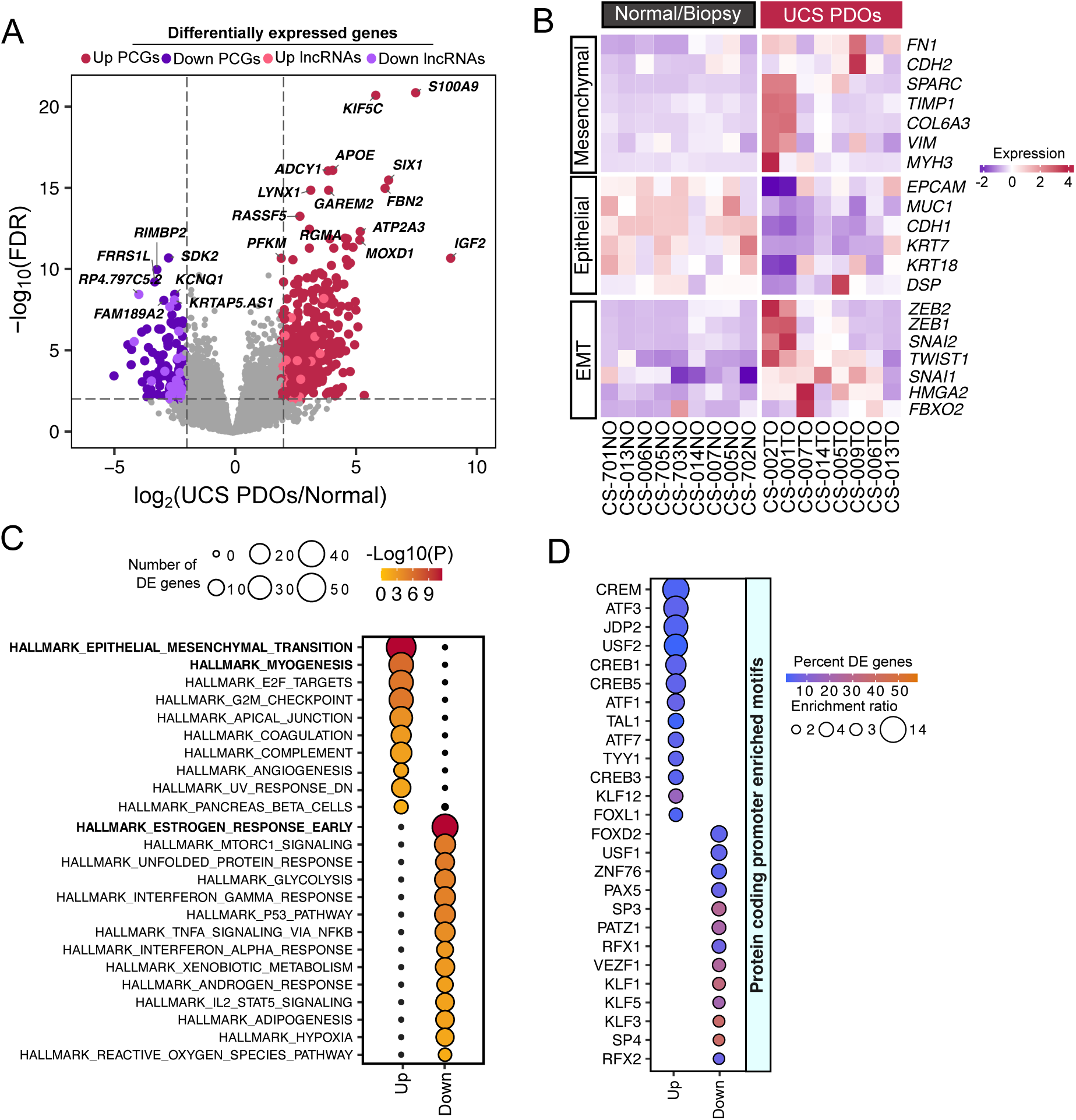
Transcriptional reprogramming in UCS PDOs reveals mesenchymal activation and CREB-associated motifs. (A) Volcano plot of differential gene expression between UCS PDOs (n = 8) and normal endometrial biopsy-derived organoids (NO; n = 9). Significantly deregulated genes (FDR < 0.05; log2 fold change≥ 1) are highlighted in red (upregulated) and blue (downregulated). Selected top upregulated and downregulated protein-coding genes and long non-coding RNAs (lncRNAs) are indicated. (B) Heatmap of lineage-associated gene signatures across UCS and normal PDOs, showing downregulation of epithelial markers and upregulation of mesenchymal and epithelial EMT-associated genes in UCS PDOs. Gene expression values are shown as Z-score-scaled log2-transformed counts. (C) Hallmark pathway enrichment analysis of differentially expressed genes comparing UCS and normal PDOs. Significance of the pathways were determined by p-values obtained using GeneSCF tool. Size of the dots represent number of differentially expressed genes in UCS involved in corresponding pathways and the color gradient shows -log10 transformed p-values, -log10(p-value). Higher the -log10(p-value) means the pathways are significantly enriched. (D) Promoter motif enrichment analysis of upregulated and downregulated protein coding genes in UCS PDOs relative to NO controls. Motif enrichment was assessed using SEA (Simple Enrichment Analysis) from MEME Suite, revealing significant enrichment of CREB-family transcription factor binding motifs (CREB1, CREB3, CREB5, CREM). Enrichment ratio was calculated using ratio of true positives and false positive sequences determined by SEA. The color gradient represents percentage of differentially expressed genes in UCS regulated by corresponding transcription factors.

Heatmap analysis of lineage-associated gene signatures demonstrated downregulation of canonical epithelial markers (e.g., *EPCAM*, *MUC1*, *CDH1*) and immune-associated genes in UCS PDOs, alongside upregulation of mesenchymal and EMT-associated genes (e.g., *VIM*, *FN1*, *SPARC*, *ZEB1*, *SNAI2*) (**Figure 6B**, **Figure S6E**). These bulk-level changes are consistent with the E/M plasticity observed at single-cell resolution in UCS PDOs (**Figure 5A**, **C-E**) and UCS primary tissue^42^. Gene set enrichment analysis further revealed significant enrichment of pathways associated with EMT, myogenesis, E2F targets, G2/M checkpoint, mTORC1 signaling, glycolysis, and inflammatory signaling (**Figure 6C**), indicating coordinated activation of proliferative and mesenchymal programs. Metabolic reprogramming was also evident, with deregulation of lipid and fatty acid metabolism genes including *CPT1C*, *APOE*, and *APOL1* (**Figure S6E**).

To identify transcription factor associations, we performed promoter motif enrichment analysis on differentially expressed protein-coding genes. Promoters of upregulated genes were significantly enriched for CREB-family binding motifs, including CREB1, CREB3, CREB5, and CREM (**Figure 6D**), implicating CREB-dependent transcriptional activation in UCS. Consistent with this, motif analysis of deregulated lncRNAs revealed enrichment of CREB binding motifs among downregulated lncRNA promoters (**Figure S6F**), suggesting context-dependent regulatory effects across coding and non-coding transcriptional programs.

Collectively, integrated bulk transcriptomic analyses reveal that UCS PDOs are defined by mesenchymal activation, metabolic remodeling, and enrichment of CREB-associated transcriptional programs, nominating CREB signaling as a potential regulatory axis underlying UCS plasticity.

### UCS PDOs model heterogeneous drug responses and nominate candidate therapeutic vulnerabilities

Having established genomically and phenotypically faithful UCS PDOs, we asked whether they could serve as a discovery platform for therapeutic interrogation, and we present the following experiments as proof of concept. As a benchmark, we screened the standard-of-care agents carboplatin (CPT) and paclitaxel (PTX), alone and in combination, which form the backbone of treatment for aggressive gynecologic malignancies^43,44^. PTX reduced viability in multiple UCS PDO lines, whereas CPT showed more variable, donor-dependent effects (**Figure 7A**, **B**; **Figure S7A**, **B**). The CPT-PTX combination produced greater cytotoxicity than single agents in several lines (**Figure 7A**, **B**), consistent with combinatorial efficacy in high-grade serous ovarian cancer^6,43,45^ and indicating that our UCS PDO models capture inter-patient variability in chemotherapy response.

**Figure 7:**
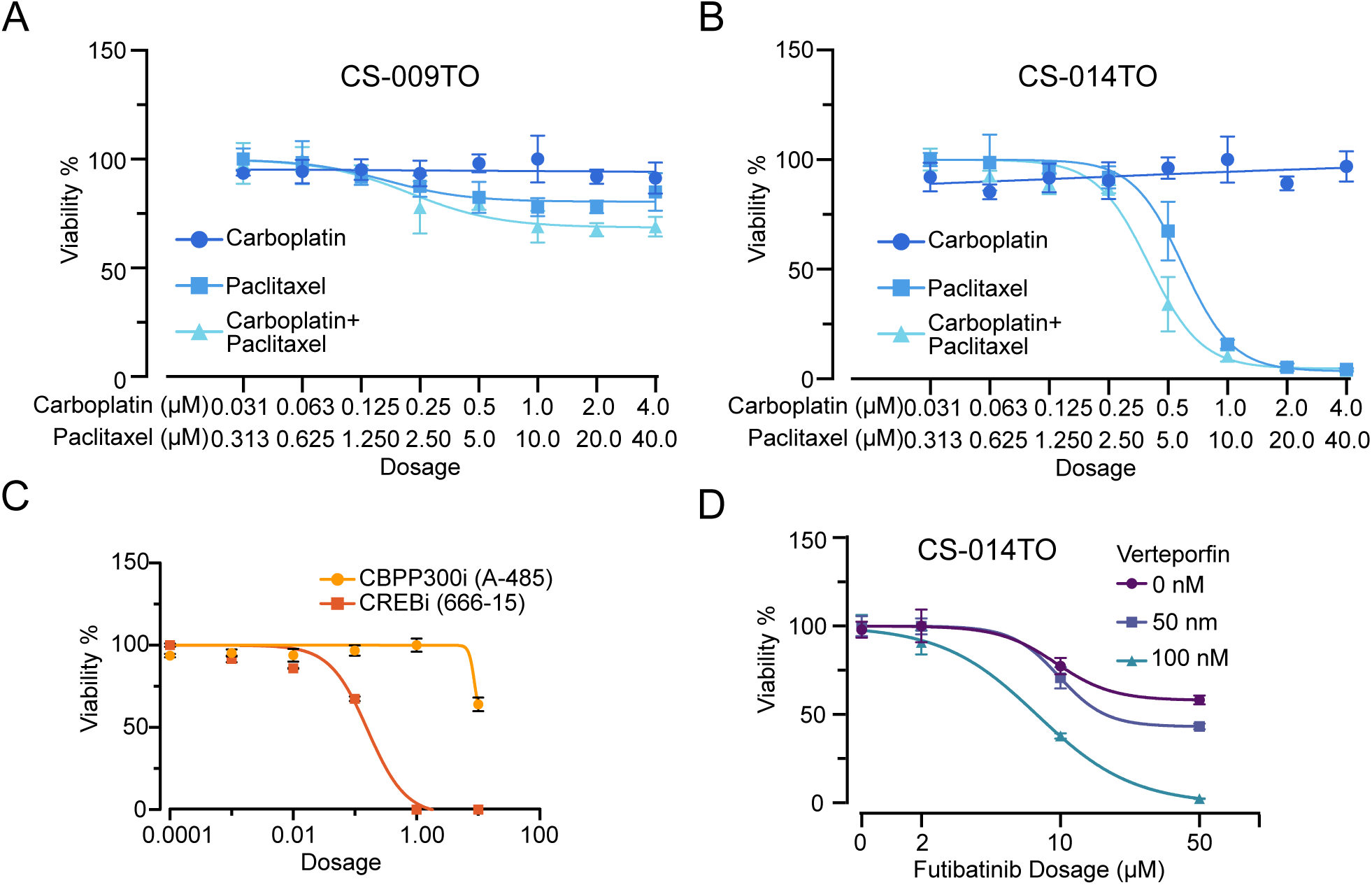
UCS PDOs serve as model to investigate in drug-response effects. (A) Dose-response curves of UCS PDO CS-009TO treated with Carboplatin (0.03125-4 µM), Paclitaxel (0.3125 µM-40 µM), and a combination of both. PDOs were treated for 6 days, and viability was quantified using the CellTiter-Glo 3D luminescent assay. Data is normalized to vehicle control and presented as mean ± SD from n = 4 technical replicates. (B) Dose-response curves of UCS PDO CS-014TO treated with Carboplatin (0.03125-4 µM), Paclitaxel (0.3125 µM-40 µM), and a combination of both. PDOs were treated for 6 days, and viability was quantified using the CellTiter-Glo 3D luminescent assay. Data is normalized to vehicle control and presented as mean ± SD from n = 4 technical replicates. (C) Dose-response curves of UCS PDO CS-014TO treated with CBPP300i (A-485, 1 nM-1000 nM) and CREBi (666-15, 1 nM-1000 nM). PDOs were treated for 6 days, and viability was quantified using the CellTiter-Glo 3D luminescent assay. Data is normalized to vehicle control and is shown as mean ± SD, with n = 8 (0 nM control) and n = 4 (treatment condition). (D) Dose-response curves of the UCS PDO line CS-014TO treated with Verteporfin (50-100 nM), Futibatinib (2-50 µM), or their combination. PDOs were treated for 6 days, and cell viability was assessed using the CellTiter-Glo 3D luminescence assay. Data is normalized to vehicle control and presented as mean ± SD, with n = 4 technical replicates for each treatment condition.

We next used the platform to test mechanism-guided hypotheses arising from our genomic and transcriptomic analyses. Motivated by the enrichment of CREB-family motifs in UCS relative to normal PDOs, we asked whether CREB signaling constitutes a candidate dependency. Pharmacologic CREB inhibition (666-15) reduced PDO viability in a dose-dependent manner, whereas inhibition of the CREB co-activator CBP/p300 (A-485) had comparatively modest effects (**Figure 7C**), nominating the CREB axis as a candidate vulnerability.

In parallel, because UCS harbors recurrent FGFR alterations in our cohort and in TCGA (**Figure 2A**, **B**), we tested FGFR inhibition with futibatinib, which alone produced only modest cytotoxicity (**Figure 7D**, **Figure S7C**). Given that adaptive activation of the Hippo effector YAP is a recurrent route of escape from FGFR inhibition and that combined FGFR and YAP blockade is effective in other cancers, including triple-negative breast, and urothelial carcinoma^46,47^, we co-targeted the two pathways. Combined futibatinib and the YAP inhibitor verteporfin reduced UCS PDO viability beyond either single agent (**Figure 7D**), consistent with a candidate cooperative FGFR-YAP dependency. While we interpret these pharmacologic results as proof of concept, they illustrate the central utility of the resource: an ancestrally inclusive, genomically faithful UCS PDO platform that recapitulates chemotherapy heterogeneity and enables systematic, hypothesis-driven discovery of cell-state-linked vulnerabilities for future mechanistic and translational studies.

## Discussion

The ability to model human disease in systems that retain its defining biological features remains a fundamental challenge in biomedical research. This is particularly critical for malignancies in which cellular identity is not fixed but dynamically regulated through transcriptional and genomic plasticity programs. UCS represents an extreme example of such complexity, yet current models do not adequately capture its biphasic architecture and underlying cell-state dynamics^23^. Here, we establish PDOs intentionally including diverse patient backgrounds that preserve both the genomic landscape and transcriptional heterogeneity of primary UCS tumors. Our UCS PDO models offer a physiologically relevant framework to interrogate disease mechanisms including cellular plasticity and therapeutic response, extending previous efforts to model UCS^23,25,26^.

Prior landmark studies established that UCS is a metaplastic carcinoma with extensive copy-number alterations, frequent *TP53* mutations and EMT transcriptional signature^10,29^. Our data demonstrate that UCS PDOs preserve these genomic properties across independent platforms. The relatively high fraction of shared variants and the strong similarity of amplification and deletion profiles indicate that the dominant tumor genome is maintained *in vitro*, even though limited unique variants suggest that some subclonal representation or culture-related selection is still possible. Our UCS models balance between the fidelity and manageable simplification of the *in vitro* culture conditions.

Single-cell analyses resolved epithelial, stem-like, and mesenchymal populations that were arranged along continuous transcriptional trajectories. Directional RNA velocity and pseudotime analyses consistently indicated transitions from epithelial toward mesenchymal-like states across independent PDO lines, although the dominant originating populations varied between donors. These observations are compatible with a model in which phenotypic diversification in UCS and other tumors arises predominantly from transcriptional state plasticity layered upon a stable genomic backbone^29,48,49^.

At the transcriptional level, UCS PDOs exhibited coordinated changes in mesenchymal, proliferative, and metabolic gene programs. The concurrent deregulation of protein-coding transcripts and long non-coding RNAs suggests broader transcriptional remodeling beyond individual pathways, consistent with emerging roles of non-coding RNAs in chromatin organization and EMT regulation^50–52^. Promoter motif analysis identified enrichment of CREB-family binding motifs, and pharmacologic CREB inhibition reduced PDO viability. CREB signaling has been previously linked to proliferation, stress adaptation, and transcriptional plasticity, and is also known to regulate non-coding RNA expression^53–57^. Together, these findings link CREB-associated transcriptional programs to cellular fitness in UCS, while further work will be required to determine whether CREB directly regulates cell state transitions or acts downstream of other signaling inputs.

Recurrent *FGFR* alterations were associated with increased transcript abundance and supported FGF-dependent PDO growth, and combined FGFR and YAP inhibition reduced viability. The observation aligns with findings in which growth factor signaling and mechanotransduction converge to reinforce EMT and stemness programs through shared transcriptional regulators^58^. Additionally, *FGFR1* activating mutations and amplifications have been reported across multiple cancer types and are known to promote tumor cell proliferation and survival, frequently correlating with adverse clinical outcomes^59,60^. Notably, prior preclinical and clinical observations in endometrial cancer, including UCS, have suggested that FGFR pathway inhibition (e.g., with pazopanib) reduces tumor burden^25,61,62^. UCS may therefore represent a signaling-integrated ecosystem in which extracellular inputs continuously reinforce transcriptional identity and plasticity. However, the extent to which these signals and pathways converge on shared transcriptional regulators remains to be defined.

While these findings establish our UCS PDOs as a robust platform to model UCS-intrinsic transcriptional plasticity, they do not fully recapitulate the multicellular complexity of the tumor microenvironment. In particular, stromal, immune, and vascular compartments, known to critically shape epithelial cell states and therapeutic response^63–65^, are not represented in the current system. Recently, we established an autologous co-culture platform using high-grade endometrial cancer PDOs and matched immune cells from the same patients for modeling epithelial-immune interactions and testing the safety and efficacy of immunotherapy drugs *in vitro*^66^. Building on these approaches, future studies should integrate UCS PDOs into more complex co-culture or multi-compartment systems to dissect how extrinsic signals reinforce or constrain transcriptional plasticity and therapy response. Additionally, therapeutic vulnerabilities, including CREB and combined FGFR-YAP inhibition, were tested in a limited number of PDO lines and require further validation, broader dose-response testing, and orthogonal assays to establish causality. In addition, E/M trajectories were inferred computationally and will require lineage tracing or live-cell approaches to confirm state transitions and directionality. Finally, although PDOs preserved key genomic and transcriptional features of primary tumors, their *in vivo* tumorigenicity and ability to predict patient-level treatment response remain to be tested.

Collectively, we have established a genomically benchmarked, ancestrally inclusive, and lineage-resolved PDO platform for interrogating cell-state plasticity and therapeutic vulnerability in UCS. This platform provides a preclinical model for future studies aimed at improving our understanding of UCS biology and developing novel therapeutic strategies.

## Methods

### Ethical approval

Fresh tumor and normal endometrial tissue specimens were obtained from patients undergoing hysterectomy for UCS from Northwell Health LIJ Medical Center. Institutional Review Board approval was obtained (study IRB #18-0897), and all patients provided informed consent prior to specimen collection.

### Endometrial tissue processing and PDOs generation

Isolation of endometrial cells and cultivation of PDOs was performed according to Katcher et al.^33^. In brief, tissue was washed with cold 1X PBS and minced into small (∼0.25 cm) fragments, before being transferred to 5 ml of RPMI media (Sigma Aldrich, #R8758) containing 1 mg/mL Collagenase (Sigma Aldrich, #C9407) and 10 µM Y-27632 dihydrochloride (Tocris, #1254). Tissue suspension was transferred to a rotating incubator (speed: 30 rpm, 37 °C) for 60-120 min depending on successful tissue dissociation. Supernatant was transferred and centrifuged at 300 g for 5 min at room temperature (RT). Supernatant was discarded and the pellet was resuspended in 3 mL of TrpLE Express Enzyme (1X, no phenol red, Thermo Fisher Scientific, #12604103) with 10 µM Y-27632 dihydrochloride followed by an incubation of 10-20 min at 37 °C with regular mixing. To stop the reacting 3 mL of ADMEM/F12 (Life Technologies, #312634028) was added and the cell suspension centrifuged at 300 g for 5 min at RT. Cells were embedded in a 70:30 mixture of Matrigel matrix (Corning, #356231): PDO media and seeded in 50 µL domes. After polymerization, normal or tumor endometrial PDO media was added. Media exchange was performed every 3-4 days.

### Passaging of endometrial PDOs

Growth was monitored and PDOs passaged every 7-14 days. Therefore, PDOs were collected by removing Matrigel using Cell Recovery Solution (Corning, #354253). After 30 min at 4 °C, PDO suspension was centrifuged at 300 g for 5 min at 4 °C. Supernatant was discarded and PDOs resuspended in 3 mL of TrpLE Express Enzyme, followed by an incubation for 5-15 min at 37 °C. 3 mL of ADMEM/F12 was added and cells centrifuged at 300 g for 5 min at 4 °C, before being seeded again in 70:30 Matrigel:PDO media mixture. Commonly, PDOs were passaged in a 1:2 ratio.

### Endometrial PDO drug screening assay

PDOs were dissociated and singularized using TrpLE Express Enzyme as previously described, centrifuged at 300 g for 5 minutes at 4 °C, and resuspended in culture medium for cell counting. Cells were embedded in a 70:30 mixture of Matrigel:PDO culture medium at a density of 200 cells per µL. Ten microliter Matrigel domes were plated in 48-well or 384-well plates and allowed to polymerize for at least 5 minutes at 37 °C prior to addition of pre-warmed culture medium per well. PDOs were cultured until day 10 prior to drug exposure.

Drug-containing media or vehicle controls were added, and treatments were refreshed on day 13. PDO viability was quantified using the CellTiter-Glo® 3D Luminescent Cell Viability Assay (Promega, #G9682) according to the manufacturer’s instructions on day 16. Briefly, culture media were aspirated and Matrigel domes were incubated with 100 µL CellTiter-Glo® 3D reagent for 30 minutes at 37°C in the dark. Lysates were transferred to opaque 96-well plates, and luminescence was measured using a SpectraMax plate reader. Luminescence values were normalized to vehicle-treated controls to determine relative viability. Per donor at least technical triplicates were used. The following compounds were used: Carboplatin (10 nM-10 µM, Sigma Aldrich, #C2538), Paclitaxel (10 nM-10 µM, Thermo Fisher Scientific, #P3456), Carboplatin + Paclitaxel (31.25 nM-4 µM +312.5 nM-40 µM) A-485 (1 nM-10 µM, MedChemExpress, #HY-107455), 666-15 (1 nM-10 µM), MedChemExpress, #HY-101120), Verteporfin (6.25 nM-400 nM, Sigma Aldrich, #SML0534), Futibatinib (0.625 µM-50µM, MedChemExpress, #HY-100818R).

### UCS PDOs bulk RNA-seq preparation and processing

PDOs were collected in Cell Recovery solution to dissolve Matrigel as previously described and afterwards lysed in 300 µl TRIzol (Zymo, #R2051). Then 300 µl 100 % ethanol (Fisher Scientific, #A4094) was added and the lysate transferred to a Zymo Spin Column and processed with the Direct-zol RNA protocol (Zymo, #R2051) according to manufacturer’s instruction.

RNA depletion and library preparation was performed using NEBNextⓇ Ultra II Directional RNA Library Prep Kit for IlluminaⓇ (Illumina, #E7765S/L) according to the manufacturer’s protocol. Prepared library was analyzed using the Agilent Bioanalyzer 2100 using high sensitivity DNA chips (Agilent, #5067-4626). Sequencing was performed using NextSeq500.

Adapter sequences were removed from the FASTQ files using Cutadapt (v5.1). Obtained Cleaned and trimmed paired-end reads files were aligned and per-samples read counts were quantified using STAR-2.5.1b with reference genome GRCh38. We used DESeq2 to perform differentially expression analysis between UCS PDOs and matched normals or biopsies. Significant DE genes were selected based on adjusted p-value (<0.05) and log-fold-change (±1.0) cutoffs. Obtained DE genes were subjected to functional enrichment analysis with human MSigDB hallmark gene sets (https://gsea-msigdb.org) and gene ontology (https://geneontology.org) using GeneSCF (v1.1-p3) tool^67^. The enriched gene sets are selected by p-value<0.05 cutoff.

### Endometrial PDOs and tissue immunohistochemistry / Hematoxylin & eosin staining

Endometrial tissue specimens were fixed in 10 % formalin for 24-48 hours at room temperature. Following fixation, tissues were dehydrated through a graded ethanol series, cleared in xylene, and embedded in paraffin using standard histological procedures. Paraffin blocks were sectioned at 4 µm thickness using a microtome and mounted onto glass slides.

PDOs were collected in Cell Recovery Solution as previously described. The suspension was transferred into a tube pre-coated with 0.1 % BSA in PBS and incubated on ice for 10-15 min with gentle pipetting every 2-3 min until Matrigel was fully dissolved. Samples were centrifuged at 300 g for 5 min at 4 °C, and the supernatant was carefully removed. The PDO pellet was fixed in 4 % paraformaldehyde (PFA) for 30 minutes at room temperature. Following fixation, samples were centrifuged at 300 g for 5 min, washed twice with 1X PBS, and centrifuged again under the same conditions. After removal of PBS, PDOs were resuspended in 2 % low-melting-point agarose. The suspension was gently mixed and allowed to solidify at RT for 10 min. The solidified agarose block containing PDOs was transferred into a histology cassette, stored in 1X PBS, and embedded and sections as described above. Hematoxylin & eosin (H&E) stainings were performed according to standard protocols.

For immunohistochemistry (IHC) stainings, slides were stained in Discovery Ultra automatic IHC stainer (Roche) following standard protocols. Briefly, after deparaffinization and rehydration, slides were subjected to antigen retrieval (Benchmark Ultra CC1, Roche) at 96°C for 64 minutes; primary Ab incubation was performed at 37 °C for 1h and Discovery multimer detection system (Discovery OmniMap HRP Roche) was used to detect and amplify immunosignals.

Primary antibodies used: E-cadherin (CST, #24E10), Ki67 (Spring Bioscience, #170822LVA), p53 (Thermo Fisher, #IHC-00010), EMA (Thermo Fisher, #Z2048MP).

### TCGA UCS and P1000 UCS gene expression and copy number correlation

BIC-Seq2 (v0.2.6)^68^ was run with default parameters to generate copy number log2 ratios., Additionally, tumor read depth was collected in 1KB bins and corrected for genomic GC content and mappability using fragCounter (https://github.com/mskilab/fragCounter). Corrected tumor coverage profiles, BAF, purity/ploidy estimates, and high confidence SVs were used as input to JaBbA (v1.1)^69^ default parameters were used, with the exception of rescue.all set to false, maxna set to 0.8, slack of 1000 and ism set to true.

We compared *FGFR1* and *FGFR3* expression against log-transformed copy number values generated by JaBbA and BIC-Seq2. Gene expression (TPM) data were obtained from the P1000 endometrial (carcinosarcoma) and TCGA-UCS cohorts, with copy number estimates derived from both tools for each cohort. Expression-copy number associations were assessed by fitting linear models using lm() in R and R² values were reported separately per cohort. We obtained TCGA-UCS gene expression data from the published dataset available on CBioPortal^29,70^. Similarly copy number correlation with gene expression profiles of UCS PDOs from this study was done using afore mentioned method.

### P1000 UCS tissue and PDOs whole genome sequencing

#### Library preparation

DNA was extracted from snap frozen patient tissue utilizing the Zymo Quick-DNA Miniprep kit (Zymo, #D3024) according to manufacturer’s instruction. DNA quality and concentration were assessed using the Nanodrop ND-1000 Spectrophotometer. Whole genome sequencing (WGS) libraries were prepared using the Truseq DNA PCR-free Library Preparation Kit (Illumina) in accordance with the manufacturer’s instructions. Briefly, 1 ug of DNA was sheared using a Covaris LE220 sonicator (adaptive focused acoustics). DNA fragments underwent bead-based size selection and were subsequently end-repaired, adenylated, and ligated to Illumina sequencing adapters. Final libraries were quantified using the Qubit Fluorometer (Life Technologies) or Spectromax M2 (Molecular Devices) and Fragment Analyzer (Advanced Analytical) or Agilent 2100 BioAnalyzer. Libraries were sequenced on an Illumina Novaseq6000 sequencer using 2×150bp cycles.

### Processing and Analysis (Alignment, Variant calling, Filtering, Annotation)

#### Pre-processing

The New York Genome Center somatic pipeline (v6) was used to process and align the WGS data and call variants. Sequencing reads for the tumor and normal samples are aligned to the reference genome GRCh38 using BWA-MEM (v0.7.15) (arXiv:1303.3997v2 [q-bio.GN]). NYGC’s ShortAlignmentMarking (v2.1) is used to mark short reads as unaligned. This tool is intended to remove spurious alignments resulting from contamination (e.g. saliva sample bacterial content) or from too aggressive alignments of short reads the size of BWA-MEM’s 19bp minimum seed length. These spurious alignments result in pileups in certain locations of the genome and can lead to erroneous variant calling.

GATK (v4.1.0)^71^ FixMateInformation is run to verify and fix mate-pair information, followed by Novosort (v1.03.01) markDuplicates to merge individual lane BAM files into a single BAM file per sample. Duplicates are then sorted and marked, and GATK’s base quality score recalibration (BQSR) was performed. The result of the pre-processing pipeline is a coordinate sorted BAM file for each sample.

#### Quality control

Once preprocessing is complete, we compute several alignment quality metrics such as average coverage, %mapped reads and %duplicate reads using GATK (v4.1.0) and an autocorrelation metric (adapted for WGS from Zhang et al.^72^) to check for unevenness of coverage. We also run Conpair^73^, a tool developed at NYGC to check the genetic concordance between the normal and the tumor sample and to estimate any inter-individual contamination in the samples.

#### Variant detection

The tumor and normal bam files are processed through NYGC’s variant calling pipeline which consists of MuTect2 (GATK v4.0.5.1)^74^, Strelka2 (v2.9.3)^75^ and Lancet (v1.0.7)^76^ for calling Single Nucleotide Variants (SNVs) and short Insertion-or-Deletion (Indels), SvABA (v0.2.1)^77^ for calling Indels and Structural variants (SVs), Manta (v1.4.0)^78^ and Lumpy (v0.2.13)^79^ for calling SVs and BIC-Seq2 (v0.2.6)^68^ for calling Copy-number variants (CNVs). Manta also outputs a candidate set of Indels which is provided as input to Strelka2 (following the developers recommendation, as it improves Strelka2’s sensitivity for calling indels >20nt).

#### Variant merging

Next, the calls are merged by variant type (SNVs, Multi Nucleotide Variants (MNVs), Indels and SVs). MuTect2 and Lancet call MNVs, however Strelka2 does not, and it also does not provide any phasing information. So to merge such variants across callers, we first split the MNVs called by MuTect2 and Lancet to SNVs, and then merge the SNV callsets across the different callers. If the caller support for each SNV in a MNV is the same, we merge them back to MNVs. Otherwise those are represented as individual SNVs in the final callset. Lancet and MantaSV are the only tools that can call deletion-insertion (delins or COMPLEX) events. Other tools may represent the same event as separate yet adjacent indel and/or SNV variants. Such events are relatively less frequent, and difficult to merge. We therefore do not merge COMPLEX calls with SNVs and Indels calls from other callers. The SVs are converted to bedpe format, all SVs below 500bp are excluded and the rest are merged across callers using bedtools^80^ pairtopair (slop of 300bp, same strand orientation, and 50% reciprocal overlap).

#### Somatic variant annotation (SNVs, Indels, CNVs, and SVs)

SNVs and Indels are annotated with Ensembl as well as databases such as COSMIC (v86)^81^, 1000Genomes (Phase3)^82^, ClinVar (201706)^83^, PolyPhen (v2.2.2)^84^, SIFT (v5.2.2)^85^, FATHMM (v2.1)^86^, gnomAD (r2.0.1)^87^ and dbSNP (v150)^88^ using Variant Effect Predictor (v93.2)^89^. For CNVs, segments with log2 > 0.2 are categorized as amplifications, and segments with log2 < - 0.235 are categorized as deletions (corresponding to a single copy change at 30% purity in a diploid genome, or a 15% Variant Allele Fraction). CNVs of size less than 20Mb are denoted as focal and the rest are considered large-scale.

We use bedtools^80^ for annotating SVs and CNVs. All predicted CNVs are annotated with germline variants by overlapping with known variants in 1000 Genomes and Database of Genomic Variants (DGV) (22). Cancer-specific annotation includes overlap with genes from Ensembl^90^ and Cancer Gene Census in COSMIC, and potential effect on gene structure (e.g. disruptive, intronic, intergenic). If a predicted SV disrupts two genes and strand orientations are compatible, the SV is annotated as a putative gene fusion candidate. Note that we do not check reading frame at this point. Further annotations include sequence features within breakpoint flanking regions, e.g. mappability, simple repeat content and segmental duplications.

### Somatic Variant Filtering

#### Panel of normals (PON)

The Panel Of Normals (PON) filtering removes recurrent technical artifacts from the somatic variant callset^74^.

#### PON generation

The Panel of Normals for SNVs, indels and SVs was created with whole-genome sequencing data from normal samples from 242 unrelated individuals. Of these, sequencing data for 148 individuals was obtained from the Illumina Polaris project which was sequenced on the HiSeqX2 platform with PCR-free sample preparation. The remaining samples were sequenced by the NYGC. Of these, 73 individuals were sequenced on HiSeqX, 11 on NovaSeq, and 10 were sequenced on both. We ran MuTect2 in artifact detection mode and Lumpy in single sample mode on these samples. For SNVs and indels, we created a PON list file with sites that were seen in two or more individuals.

For SVs, we used SURVIVOR (v1.0.3)^91^ to merge Lumpy calls. Variants were merged if they were of the same type, had the same strand orientation, and were within 300bp of each other (maximum distance). We did not specify a minimum size. After merging SVs, we used these calls as a PON list.

#### PON filtering

For SNVs and Indels, we use the PON list to filter the somatic variants in the merged SNV and indel files. To filter our somatic SV callset, we merge our PON list with our callset using bedtools pairtopair (slop of 300bp, same strand orientation, and 50% reciprocal overlap), and filtered those SVs found in two or more individuals in our PON.

#### Common germline variants

In addition to the PON filtering, we remove SNVs and Indels that have minor allele frequency (MAF) of 1% or higher in either 1000Genomes (phase 3) or gnomAD (r2.0.1)^87^, and SVs that overlap DGV and 1000Genomes (phase3). CNVs are annotated with DGV and 1000 Genomes but not filtered.

#### All somatic and high-confidence variants

Variants that pass all of the above-mentioned filters are included in our final somatic callset (hereby referred to as AllSomatic). For SNVs, indels and SVs, we also annotate a subset of the somatic callset as high confidence. For SNVs and indels, high confidence calls are defined as those that are either called by two or more variant callers or called by one caller and also seen in the Lancet validation calls or in the Manta SV calls.

For structural variants, high confidence calls are taken from the somatic callset if they meet the following criteria: called by 2 or more variant callers or called by Manta or Lumpy with either additional support from nearby CNV changepoint or split-read support from SplazerS^87^, an independent tool used to calculate the number of split-reads supporting SV breakpoints. An SV is considered supported by SplazerS if it found at least 3 split-reads in the tumor only. Nearby CNV changepoints are determined by overlapping BIC-Seq2 calls with the SV callset using bedtools closest. An SV is supported by a CNV changepoint if the breakpoint of the CNV is within 1000bp of an SV breakpoint.

### MSI detection

We run MANTIS (v1.0.4)^32^ for Microsatellite Instability (MSI) detection in microsatellite loci (found using RepeatFinder, a tool included with MANTIS). A sample is considered to be 6 microsatellite unstable if it’s Step-Wise Difference score reported by MANTIS is greater than 0.4 (or 0.62 in absence of a matched-normal). Otherwise, it is microsatellite stable3 (MSS).

### Genetic ancestry estimation from whole genome

Ancestry proportion is determined by the software ADMIXTURE v1.3.0^30^, which uses a maximum likelihood-based method to estimate the proportion of reference population ancestries in a sample. We genotyped the reference markers generated from 1,964 unrelated 1000 Genomes project samples directly on the samples using GATK pileup. Individuals from populations MXL (Mexican Ancestry from Los Angeles USA), ACB (African Caribbean in Barbados), and ASW (African Ancestry in Southwest US) were excluded from the reference due to being putatively admixed. The reference was further filtered by using only SNP markers with a minimum minor allele frequency (MAF) of 0.01 overall and 0.05 in at least one 1000 genomes superpopulation. Variants are additionally linkage disequilibrium (LD) pruned using PLINK v1.9 with a window size of 500kb, a step size of 250kb and r2 threshold of 0.2. The analysis results in a proportional breakdown of each sample into 5 continental populations (AFR, AMR, EAS, EUR, SAS) and 23 sub-populations.

### UCS panel sequencing processing and variant calling

CSHL Next Generation Sequencing Core Facility performed capture based targeted gene panel sequencing (for a panel of potential cancer driver genes). Briefly, we initially used a panel of 143 cancer genes (based on the hg19 reference genome) with a total of ∼4000 probes for capture. We later updated to a new panel of 163 genes (including all original genes) using ∼7700 Twist probes based on the hg38 reference genome. The captured DNA was prepared into libraries using the Twist Library Preparation EF Kit 2.0 and Twist UDI TruSeq compatible adapters. Final captured libraries were sequenced on an Illumina NextSeq500 using paired-end 150bp reads to a coverage of ∼100–300x.

### UCS whole exome (WE) sequencing processing and variant calling

CSHL Next Generation Sequencing Core Facility prepared libraries with the KAPA HyperPlus Library Preparation kit. Briefly, gDNA was enzymatically fragmented in a thermal cycler for 30 minutes, followed by end repair and A-tailing. Illumina compatible dual indexes were ligated and libraries were amplified with 7 cycles of pcr enrichment, quantified by qubit and pooled equimolar. Following manufacturer protocol, libraries were hybridized to NimbleGen SeqCap EZ Library v3.0 probes^92^ for 72 hours in a thermal cycler. Post hybridization, capture reactions were recovered and washed, followed by 12 cycles of post capture pcr enrichment. These post capture libraries were purified, and quantified with qubit and qpcr. Agilent tapestation was to determine insert fragment length. Post capture libraries were pooled and sequenced on an Illumina NextSeq500 PE150 high output run to ∼75-100X coverage.

Whole exome sequences were aligned with Burrows-Wheeler Aligner (BWA) v0.7.17^93^ (RRID:SCR_010910) to the human genome (GRCh38.p13 assembly). Picard v2.21.8 (http://broadinstitute.github.io/picard/) was used to removed duplicated sequences. A recalibration step was done with GATK v4.1.2.0^71^.

Copy number variants (CNV) were identified with CNVkit v0.9.6^94^. The results were post-processed with CNprep^95^ package v2.1.1 to calculate the copy number values relative to their central values for each segment (median deviation in log2ratio). Thresholds to assign a significant alteration were set to p-value < 0.001 for deletions and p-value > 0.999 for amplifications. The chromosomes Y and M have been removed from the analyses.

### Mutational analysis

The sequencing reads were aligned to the hg19 or hg38 reference genome (depending upon capture probes used) using BWA^96^, followed by BAM formatting and sorting with Samtools^97^. PCR duplicates were removed with Picard (https://broadinstitute.github.io/picard/), and Bamtools^98^ was used to select a minimum mapping quality of 20 and proper read pairing. Coverage of the target regions was assessed using Picard HSmetrics to ensure adequate coverage for confident variant detection. Variants were called using VarScan2^99^ in somatic mode to stratify germline versus somatic variants. Resulting variants were annotated with Annovar^100^ to cover a broad range of prediction tools. We selected rare loss of function variants (nonsense, frameshift, splice site) with frequency less than 1% in the Gnomad, ExAC, EVS and 1000 Genomes databases. Missense and in-frame indel variants were selected if they were noted as pathogenic by ClinVar^101^, or if they were both rare and present in COSMIC^81^, or if they are both rare and found to be present in the TCGA cohorts.

In order to harmonize variants between captures, we used the liftOver tool from the UCSC Genome Browser to convert coordinates form hg19 to hg38.

Oncoplots are generated from these candidate variants using Maftools^102^.

### UCS whole genome (SMASH) sequencing processing

SMASH sequencing data were processed using a custom SMASH Analytics pipeline developed in-house, as previously described^103^, with implementation available at https://github.com/docpaa/smash-paper. Sequencing reads were aligned to the human reference genome (hg19 assembly).

Copy number variants (CNV) were detected using CNprep^95^ package v2.1.1 to obtain the median deviation in log2ratio. Thresholds to assign a significant alteration were set to p-value < 0.001 for deletions and p-value > 0.999 for amplifications. The chromosomes Y and M have been removed from the analyses. The graphical representation of the amplification and deletion events through the genome has been generated with the Bioconductor gtrellis^104^ package v1.28.0.

### UCS PDOs-tumor CNV comparison in whole exome and SMASH

Similarity between copy number profiles has been calculated with Bioconductor CNVMetrics^105,106^ package v1.6.0. The copy number profiles, deletion and amplification separated in different sets, were used as input. For each pair of profiles, the Sørensen coefficient is used as the similarity measure. The Sørensen coefficient is obtained by dividing twice the size of the intersection of the amplified/deleted regions by the sum of the size of the amplified/deleted regions for the two profiles. For each pair of profiles, 500 synthetic profiles were generated for one profile of the pair (when a metric could not be successfully calculated with a synthetic profile, the synthetic profile was discarded). The p-value was obtained by calculating the ratio of synthetic profiles with equal or higher coefficient that the one obtained with the real pair of profiles (one observation is always added to the calculation).

The similarity heatmaps were generated with Bioconductor ComplexHeatmap^104^ package v2.12.0 using the Euclidean distance and the complete method for clustering.

### Analyzing somatic variants and ancestry using scRNA-seq from UCS PDOs

First, FASTQ files were extracted from scRNA-seq alignment done on cellranger from 10X Chromium using GRCh38 2016 release 84 reference and converted BAM files to FASTQ with the help of 10X bamtofastq software version 1.4.1 (https://support.10xgenomics.com/docs/bamtofastq). The FASTQ files were used as input to generate new alignments on 10X Chromium GRCh38 2020A reference with 10X Chromium cellranger software version 7.0.0^107^. Those alignments are used for further processing. Variant calling was done following the GATK RNA-seq Best Practices^71^ except for the duplicate removal step which was skipped. The first steps of the pipeline were run with GATK software version 4.2.5.0, and the final output was the recalibrated BAM files. At last, Picard software version 2.26.11 (https://broadinstitute.github.io/picard/) was used to resort the BAM files according to GENCODE version 39 genome order^108^. Somatic variants were extracted for each cell with both SCExecute software version 1.3.2^109^ and snp-pileup tool from FACETS software version 0.6.1 (Shen et al 2016). The list of somatic variants used as input was specific to each PDO samples as obtained for the variant calling done on the exome sequencing.

Genetic ancestry on scRNA-seq was done with RAIDS software v0.99.15 following the pipeline presented in Belleau et al. 2023^110^ with the inference step changed from principal-component analysis to admixture analysis with ADMIXTURE software version 1.3.0^30^. The list of variants used as input was obtained with snp-pileup tool from FACETS software version 0.6.1^111^ run on the recalibrated BAM files directly.

### UCS single cell profiling by Chromium 10X sequencing and analysis

For cell isolation from endometrial tissue, samples were processed as described above for endometrial tissue processing and PDOs generation. Following termination of trypsin digestion and centrifugation, cell pellets were resuspended in 3 mL of fresh ADMEM/F12 and filtered through a 40 µm cell strainer (Corning, #431750). Dead cells were subsequently removed using the Dead Cell Removal Kit (Miltenyi Biotec, #130-090-101) according to the manufacturer’s instructions using a MACS separation system. Final cell suspensions exhibited 80-90% viability with yields of at least1.5 million cells.

PDO lines maintained for at least two passages after thawing or initial establishment were harvested for single-cell preparation. Matrigel was dissolved by incubating PDOs in Cell Recovery Solution for 1 h at 4 °C. PDOs were washed and enzymatically dissociated using TrypLE Express for 10-45 min, with gentle mechanical trituration by pipetting every 5 min to promote generation of single cells. Digestion was monitored by brightfield microscopy and quenched by dilution with serum-free culture medium once an adequate fraction of single cells was observed. Cell suspensions were filtered through a 40 µm strainer, pelleted, and resuspended in ADMEM/F12. Dead cells were removed using the Dead Cell Removal Kit according to the manufacturer’s instructions. Final preparations typically yielded in over 1.5 million cells with 80-90% viability. Samples are processed for sequencing with Chromium Next GEM Chip G (10x Genomics). GEM generation and barcoding, reverse transcription, cDNA generation and library construction using 3ʹ Gene Expression Library Construction using the Chromium Single Cell 30 Library (v3 chemistry) following the manufacturer’s protocol. Dual-indexed, single-cell libraries were pooled and sequenced in paired-end reads on Novaseq (Illumina).

#### Single-cell analysis

Raw reads from FASTQ files are aligned to reference genome GRCh38 and quantified for GRCh38.84 gene annotation using CellRanger (v.6.0.0) (https://10xgenomics.com). First, the CellRanger *mkfastq* command with the CellRanger sample sheet was used to demultiplex the base call files for each flow cell into FASTQ files. Second, the CellRanger *count* command was called to generate single cell feature counts for each library by specifying the library name in the argument. The filtered feature barcode matrix was used for further data analysis.

#### Quality check and clustering

The downstream analysis using feature barcode matrix was performed using Seurat (v.3.0.0) package. Individual patients with their feature barcode matrix libraries were converted into Seurat object using *Read10X* and *CreateSeuratObject*. For each individual patients we removed cells with high mitochondrial content (>20%) and cells with less than 200 genes. These cut-offs are based on QC inspection and previous single cell studies on endometrium^112,36,95^. Subsequently, the gene in remaining 97,622 cells were log normalized, variable genes detected, scaled and the principal components computed. The top principal components were identified and used for the UMAP and TSNE dimensionality reduction. Normal endometrium singe cell data from Wang et al.^112^, was filtered using the cutoffs mentioned in the publication and the clustering is performed like CS patients.

#### Heterogeneity and assignment of cell types

Initial clustering was performed, and CS patients showed highly heterogenous profiles. Individual clusters are assigned to a patient based on proportion of cells from the patient contributing to the cluster. Hence on the gene markers are predicted per cluster in individual CS patient by *FindAllMarkers* function from Seurat. This method of assigning gene markers and cell types preserved both molecular and cellular-level heterogeneity in CS patients. Similar approach was used in prediction of gene markers and cell type assignment for normal endometrium singe cell data from Wang et al. Scoring of cells for different functional modules are done using *AddModuleScore_UCell* function from UCell R package.

### UCS single cell differential expression analysis

Differential expression (DE) analysis between normal endometrium and CS predicted cancer cells were performed by implementing Libra using *run_de* function (https://github.com/neurorestore/Libra)^113^. We used single cell appropriate statistical method Wilcoxon Rank-Sum test to find differentially expressed genes. Similarly, DE analysis was performed between CS primary and metastatic stages, between normal endometrium immune profile and CS tumor infiltrating immune profile. Significant DE genes were selected based on adjusted p-value (<0.05) and log-fold-change (±1.0) cutoffs. Obtained DE genes were subjected to functional enrichment analysis with human MSigDB hallmark gene sets (https://gsea-msigdb.org) and gene ontology (https://geneontology.org) using GeneSCF (v1.1-p3) tool^67^. The enriched gene sets are selected by p-value<0.05 cutoff.

### UCS promoter sequence based motif enrichment analysis

Using custom scripts we extracted nucleotide sequences (FASTA) from the promoters (± 250 bp from TSS) of list of significantly differentially expressed genes between CS predicted cancer cells and normal endometrium (https://github.com/decodebiology/extract_promoter_sequence) with GRCh38.84 gene and genome annotation. Obtained FASTA sequences were used to find enriched transcription factor motifs using SEA (v5.5.5) from MEME tool. Motif database HOCOMOCO (v11) full human was used, and the shuffled primary sequences preserving 3-mer frequencies are considered as control sequences. Also, 10% of the input sequences were randomly assigned to the hold-out set to improve p-value accuracy. The motifs are selected based on enrichment E-value of 10 or smaller. Motifs analysis was performed independently on both up and downregulated genes.

## Data and Code Availability

The bulk RNA-seq and single-cell RNA-seq data of patient-derived organoids (PDOs) has been deposited in GEO with accessions GSE307568 and GSE326304 respectively. Single-cell RNA-seq datasets from patient tissues can be accessible from GEO using accession, GSE326307. The P1000 cohort of UCS genomic sequencing data and the genomic data generated in this study will be deposited in dbGAP repository. All the pipelines and the code used for the analysis has been deposited in GitHub, https://github.com/decodebiology/Carcinosarcoma_organoids/. The single-cell dataset generated and/or analyzed in this study can be explored through an interactive portal available at: https://decodebiology.shinyapps.io/UCS_PDOs/.

## Acknowledgements

We thank the members of the Beyaz lab for critical discussions. We thank Northwell Health Biospecimen Repository and the Gynecologic Oncology staff for assistance in recruitment of patients from diverse demographics and acquisition of specimens for this study. We thank Cold Spring Harbor Laboratory Cancer Center Shared Resources (Flow Cytometry, Microscopy, Sequencing, Organoid and Histology Core Facilities) supported in part by the National Cancer Institute Cancer Center Support Grant (5P30CA045508). This work was financially supported by grants to S.B. from the National Cancer Institute (R37CA292807), Oliver S. and Jennie R. Donaldson Charitable Trust, the Mark Foundation for Cancer Research (20-028-EDV), the Beverly Hazelkorn Uterine Cancer Research Foundation, the Cold Spring Harbor Laboratory and Northwell Health Affiliation, New York Genome Center Polyethnic-1000 Initiative. This work was performed with assistance from the US National Institutes of Health Grant S10OD028632-01. The results shown here are in part based upon data generated by the TCGA Research Network: https://www.cancer.gov/tcga. P.B. is supported by the National Cancer Institute’s Informatics Technology for Cancer Research (ITCR) program Grant U01CA289357. The sample processing, computations, and depositions of the datasets from this study and TCGA was performed on High Performance Computing (HPC) resources Elzar and Agastya provided by Cold Spring Harbor Laboratory and Indian Institute of Technology Jammu respectively.

## References

1. Bogani, G. et al. Endometrial carcinosarcoma. Int. J. Gynecol. Cancer 33, 147–174 (2023).

2. Maiorano, M. F., Cormio, G., Maiorano, B. A. & Loizzi, V. Uterine Carcinosarcoma (UCS): A Literature Review and Survival Analysis from a Retrospective Cohort Study. Cancers 16, 3905 (2024).

3. Gopinatha Pillai, M. S., et al. Uterine carcinosarcoma: Unraveling the role of epithelial-to-mesenchymal transition in progression and therapeutic potential. FASEB J. 38, e70132 (2024).

4. Beliaeva, A. & Bakhidze, E. Diagnosis and treatment of uterine carcinosarcoma. J. Clin. Oncol. 38, e18111–e18111 (2020).

5. Tung, H.-J. et al. Management and Prognosis of Patients with Recurrent or Persistent/Progressive Uterine Carcinosarcoma. Curr. Oncol. 29, 7607–7623 (2022).

6. Powell, M. A. et al. Randomized Phase III Trial of Paclitaxel and Carboplatin Versus Paclitaxel and Ifosfamide in Patients With Carcinosarcoma of the Uterus or Ovary: An NRG Oncology Trial. J. Clin. Oncol. 40, 968–977 (2022).

7. Mahdi, H. et al. Evolving treatment paradigms in metastatic or recurrent low-grade endometrial cancer: When is hormonal-based therapy the preferred option? Int. J. Gynecol. Cancer Off. J. Int. Gynecol. Cancer Soc. 33, 1675–1681 (2023).

8. Pham, E. N. B. et al. Quality of life and survival in patients with uterine carcinosarcoma: A tertiary center observational study. Gynecol. Oncol. Rep. 57, 101679 (2025).

9. Pham, E. N. B. et al. Molecular diversity in uterine carcinosarcoma: Beyond TP53. Gynecol. Oncol. 198, 75–83 (2025).

10. Gotoh, O. et al. Clinically relevant molecular subtypes and genomic alteration-independent differentiation in gynecologic carcinosarcoma. Nat. Commun. 10, 4965 (2019).

11. Kandoth, C. et al. Integrated genomic characterization of endometrial carcinoma. Nature 497, 67–73 (2013).

12. Lee, M.-K. et al. Cumulative abdominal obesity exposure and progressive risk of endometrial cancer in young women: a nationwide cohort study. Int. J. Obes. 49, 2094–2101 (2025).

13. Liao, C.-I. et al. Increasing incidence of uterine carcinosarcoma: A United States Cancer Statistics study. Gynecol. Oncol. Rep. 40, 100936 (2022).

14. Abel, M. K. et al. Racial disparities in high-risk uterine cancer histologic subtypes: A United States Cancer Statistics study. Gynecol. Oncol. 161, 470–476 (2021).

15. Ben-David, U. et al. Genetic and transcriptional evolution alters cancer cell line drug response. Nature 560, 325–330 (2018).

16. Collins, A. et al. Patient-derived explants, xenografts and organoids: 3-dimensional patient-relevant pre-clinical models in endometrial cancer. Gynecol. Oncol. 156, 251–259 (2020).

17. Maenhoudt, N., De Moor, A. & Vankelecom, H. Modeling Endometrium Biology and Disease. J. Pers. Med. 12, (2022).

18. Xue, Y. et al. Preclinical research models for endometrial cancer: development and selection of animal models. Front. Oncol. 15, 1512616 (2025).

19. Clevers, H. Modeling Development and Disease with Organoids. Cell 165, 1586–1597 (2016).

20. Maru, Y. et al. Establishment and characterization of multiple patient-derived organoids from a case of advanced endometrial cancer. Hum. Cell 37, 840–853 (2024).

21. Dahl, M. J. et al. Gemcitabine combination therapies induce apoptosis in uterine carcinosarcoma patient-derived organoids. Front. Oncol. 14, 1368592 (2024).

22. Moufarrij, S. et al. TROP2 as a novel target in uterine carcinosarcoma organoid models. Gynecol. Oncol. 190, S204 (2024).

23. Cuppens, T. et al. Establishment and characterization of uterine sarcoma and carcinosarcoma patient-derived xenograft models. Gynecol. Oncol. 146, 538–545 (2017).

24. Moufarrij, S. et al. TROP2 expression and therapeutic targeting in uterine carcinosarcoma. Gynecol. Oncol. 197, 129–138 (2025).

25. Wong, N. K. Y. et al. Establishment and characterization of preclinical models of human gynecologic tract carcinosarcomas demonstrates targetable FGFR1 alterations. Transl. Oncol. 63, 102591 (2026).

26. Maru, Y. et al. Establishment and characterization of multiple patient-derived organoids from a case of advanced endometrial cancer. Hum. Cell 37, 840–853 (2024).

27. Park, A. B. et al. Racial disparities in survival among women with endometrial cancer in an equal access system. Gynecol. Oncol. 163, 125–129 (2021).

28. Long, B., Liu, F. W. & Bristow, R. E. Disparities in uterine cancer epidemiology, treatment, and survival among African Americans in the United States. Gynecol. Oncol. 130, 652–659 (2013).

29. Cherniack, A. D. et al. Integrated Molecular Characterization of Uterine Carcinosarcoma. Cancer Cell 31, 411–423 (2017).

30. Alexander, D. H., Novembre, J. & Lange, K. Fast model-based estimation of ancestry in unrelated individuals. Genome Res. 19, 1655–1664 (2009).

31. Maiorano, M. F. P., Cormio, G., Maiorano, B. A. & Loizzi, V. Uterine Carcinosarcoma (UCS): A Literature Review and Survival Analysis from a Retrospective Cohort Study. Cancers 16, (2024).

32. Kautto, E. A. et al. Performance evaluation for rapid detection of pan-cancer microsatellite instability with MANTIS. Oncotarget 8, 7452–7463 (2017).

33. Katcher, A. et al. Establishing patient-derived organoids from human endometrial cancer and normal endometrium. Front. Endocrinol. 14, 1059228 (2023).

34. Barbi, M. et al. Generation and Maintenance of Patient-Derived Endometrial Cancer Organoids. Bio-Protoc. 14, e5093 (2024).

35. Boretto, M. et al. Patient-derived organoids from endometrial disease capture clinical heterogeneity and are amenable to drug screening. Nat. Cell Biol. 21, 1041–1051 (2019).

36. Ornitz, D. M. & Itoh, N. The Fibroblast Growth Factor signaling pathway. Wiley Interdiscip. Rev. Dev. Biol. 4, 215–266 (2015).

37. ElHarouni, D., Al-Jazrawe, M., Choi, S., Dede, M., Hinoue, T., Misek, S. A., Noh, H., Zanella, L., Tseng, Y.-Y., Francies, H. E., Plenker, D., Kyi, C. W., Perez-Mayoral, J., Stine, M. J., Tonsing-Carter, E., Agarwal, R., Zenklusen, J. C., Clinton, J. M., Shelton, J. M., Chu, T. R., Hooper, W. F., Loinaz, X., Keskula, P., Lee, J. A., Kuhlers, P. C., Tercan, B., Boj, S. F., Vasciaveo, A., Tomassoni, L., Crawford, J. M., Walsh, S., Sinai, C., Bhatia, S., Sridevi, P., Patel, H., Cerone, M. A., The HCMI Network, Ellrott, K., Kuo, C. J., Elemento, O., Beyaz, S., Corbo, V., Spector, D. L., Beroukhim, R., Ferguson, M. L., Cherniack, A. D., Laird, P. W., Robine, N., McPherson, A., Hoadley, K. A., Garnett, M. J., Tuveson, D. A., Califano, A., Spellman, P. T., Ligon, K. L., Gerhard, D. S., Staudt, L. M., Boehm, J. S. A compendium of next-generation patient-derived models for diverse cancers.

38. Murali, R. et al. High-grade Endometrial Carcinomas: Morphologic and Immunohistochemical Features, Diagnostic Challenges and Recommendations. Int. J. Gynecol. Pathol. Off. J. Int. Soc. Gynecol. Pathol. 38 **Suppl 1**, S40–S63 (2019).

39. Zhao, S. et al. Mutational landscape of uterine and ovarian carcinosarcomas implicates histone genes in epithelial-mesenchymal transition. Proc. Natl. Acad. Sci. U. S. A. 113, 12238–12243 (2016).

40. Zakrzewski, P. K. Canonical TGFβ Signaling and Its Contribution to Endometrial Cancer Development and Progression-Underestimated Target of Anticancer Strategies. J. Clin. Med. 10, (2021).

41. Alexander, D. H., Novembre, J. & Lange, K. Fast model-based estimation of ancestry in unrelated individuals. Genome Res. 19, 1655–1664 (2009).

42. Subhash, S. et al. A Single-Cell Atlas of Uterine Carcinosarcoma from Diverse Ancestries. bioRxiv 2026.04.21.720013 (2026) doi:10.64898/2026.04.21.720013.

43. Chan John K., et al. Weekly vs. Every-3-Week Paclitaxel and Carboplatin for Ovarian Cancer. N. Engl. J. Med. 374, 738–748.

44. Miller, D. S. et al. Carboplatin and Paclitaxel for Advanced Endometrial Cancer: Final Overall Survival and Adverse Event Analysis of a Phase III Trial (NRG Oncology/GOG0209). J. Clin. Oncol. Off. J. Am. Soc. Clin. Oncol. 38, 3841–3850 (2020).

45. Lee, M. X. & Tan, D. S. Weekly versus 3-weekly paclitaxel in combination with carboplatin in advanced ovarian cancer: which is the optimal adjuvant chemotherapy regimen? J. Gynecol. Oncol. 29, e96 (2018).

46. Pettitt, G. A. et al. Development of resistance to FGFR inhibition in urothelial carcinoma via multiple pathways in vitro. J. Pathol. 259, 220–232 (2023).

47. Li, Y. et al. FGFR-inhibitor-mediated dismissal of SWI/SNF complexes from YAP-dependent enhancers induces adaptive therapeutic resistance. Nat. Cell Biol. 23, 1187–1198 (2021).

48. Quintanal-Villalonga, Á. et al. Lineage plasticity in cancer: a shared pathway of therapeutic resistance. Nat. Rev. Clin. Oncol. 17, 360–371 (2020).

49. Pastushenko, I. & Blanpain, C. EMT Transition States during Tumor Progression and Metastasis. Trends Cell Biol. 29, 212–226 (2019).

50. Khanbabaei, H. et al. Non-coding RNAs and epithelial mesenchymal transition in cancer: molecular mechanisms and clinical implications. J. Exp. Clin. Cancer Res. CR 41, 278 (2022).

51. Han, P. & Chang, C.-P. Long non-coding RNA and chromatin remodeling. RNA Biol. 12, 1094–1098 (2015).

52. Mitra, R. et al. Decoding critical long non-coding RNA in ovarian cancer epithelial-to-mesenchymal transition. Nat. Commun. 8, 1604 (2017).

53. Mehra, S., Singh, S. & Nagathihalli, N. Emerging Role of CREB in Epithelial to Mesenchymal Plasticity of Pancreatic Cancer. Front. Oncol. 12, 925687 (2022).

54. Yin, Y. et al. The CRTC-CREB axis functions as a transcriptional sensor to protect against proteotoxic stress in Drosophila. Cell Death Dis. 13, 688 (2022).

55. Hong, J. et al. cAMP response element–binding protein: A credible cancer drug target. J. Pharmacol. Exp. Ther. 392, (2025).

56. Wang, J. et al. CREB up-regulates long non-coding RNA, HULC expression through interaction with microRNA-372 in liver cancer. Nucleic Acids Res. 38, 5366–5383 (2010).

57. Jiang, L.-Y. et al. CREB-induced LINC00473 promotes chemoresistance to TMZ in glioblastoma by regulating O6-methylguanine-DNA-methyltransferase expression via CEBPα binding. Neuropharmacology 243, 109790 (2024).

58. Ilyas, S. I. et al. A Hippo and Fibroblast Growth Factor Receptor Autocrine Pathway in Cholangiocarcinoma. J. Biol. Chem. 291, 8031–8047 (2016).

59. Turner, N. et al. FGFR1 amplification drives endocrine therapy resistance and is a therapeutic target in breast cancer. Cancer Res. 70, 2085–2094 (2010).

60. Dieci, M. V., Arnedos, M., Andre, F. & Soria, J. C. Fibroblast growth factor receptor inhibitors as a cancer treatment: from a biologic rationale to medical perspectives. Cancer Discov. 3, 264–279 (2013).

61. Sawayama, S. et al. Efficacy of pazopanib in FGFR1-amplified uterine carcinosarcoma: A case report. Gynecol. Oncol. Rep. 41, 100993 (2022).

62. Sengal, A. T. et al. Endometrial cancer PDX-derived organoids (PDXOs) and PDXs with FGFR2c isoform expression are sensitive to FGFR inhibition. *Npj Precis*. Oncol. 7, 127 (2023).

63. Grant, G. & Ferrer, C. M. The role of the immune tumor microenvironment in shaping metastatic dissemination, dormancy, and outgrowth. Trends Cell Biol. https://doi.org/10.1016/j.tcb.2025.05.006 doi:10.1016/j.tcb.2025.05.006.

64. Anderson, N. M. & Simon, M. C. The tumor microenvironment. Curr. Biol. CB 30, R921–R925 (2020).

65. Raghavan, S. et al. Microenvironment drives cell state, plasticity, and drug response in pancreatic cancer. Cell 184, 6119–6137.e26 (2021).

66. Chung, C. et al. Ancestrally Diverse Autologous Patient-Derived Organoid–Immune Cell Coculture Platform for Addressing Immunotherapeutic Outcome Disparities in High-Grade Endometrial Cancer. Cancer Res. Commun. 6, 657–671 (2026).

67. Subhash, S. & Kanduri, C. GeneSCF: a real-time based functional enrichment tool with support for multiple organisms. BMC Bioinformatics 17, 365 (2016).

68. Xi, R., Lee, S., Xia, Y., Kim, T.-M. & Park, P. J. Copy number analysis of whole-genome data using BIC-seq2 and its application to detection of cancer susceptibility variants. Nucleic Acids Res. 44, 6274–6286 (2016).

69. Hadi, K. et al. Distinct Classes of Complex Structural Variation Uncovered across Thousands of Cancer Genome Graphs. Cell 183, 197–210.e32 (2020).

70. Cerami, E. et al. The cBio cancer genomics portal: an open platform for exploring multidimensional cancer genomics data. Cancer Discov. 2, 401–404 (2012).

71. McKenna, A. et al. The Genome Analysis Toolkit: a MapReduce framework for analyzing next-generation DNA sequencing data. Genome Res. 20, 1297–1303 (2010).

72. Zhang, L. & Zhang, L. Use of autocorrelation scanning in DNA copy number analysis. Bioinforma. Oxf. Engl. 29, 2678–2682 (2013).

73. Bergmann, E. A., Chen, B.-J., Arora, K., Vacic, V. & Zody, M. C. Conpair: concordance and contamination estimator for matched tumor-normal pairs. Bioinforma. Oxf. Engl. 32, 3196–3198 (2016).

74. Cibulskis, K. et al. Sensitive detection of somatic point mutations in impure and heterogeneous cancer samples. Nat. Biotechnol. 31, 213–219 (2013).

75. Kim, S. et al. Strelka2: fast and accurate calling of germline and somatic variants. Nat. Methods 15, 591–594 (2018).

76. Narzisi, G. et al. Genome-wide somatic variant calling using localized colored de Bruijn graphs. *Commun*. Biol. 1, 20 (2018).

77. Wala, J. A. et al. SvABA: genome-wide detection of structural variants and indels by local assembly. Genome Res. 28, 581–591 (2018).

78. Chen, X., et al. Manta: rapid detection of structural variants and indels for germline and cancer sequencing applications. Bioinforma. Oxf. Engl. 32, 1220–1222 (2016).

79. Layer, R. M., Chiang, C., Quinlan, A. R. & Hall, I. M. LUMPY: a probabilistic framework for structural variant discovery. Genome Biol. 15, R84 (2014).

80. Quinlan, A. R. & Hall, I. M. BEDTools: a flexible suite of utilities for comparing genomic features. Bioinforma. Oxf. Engl. 26, 841–842 (2010).

81. Tate, J. G. et al. COSMIC: the Catalogue Of Somatic Mutations In Cancer. Nucleic Acids Res. 47, D941–D947 (2019).

82. Auton, A. et al. A global reference for human genetic variation. Nature 526, 68–74 (2015).

83. Landrum, M. J. et al. ClinVar: public archive of relationships among sequence variation and human phenotype. Nucleic Acids Res. 42, D980–985 (2014).

84. Adzhubei, I., Jordan, D. M. & Sunyaev, S. R. Predicting functional effect of human missense mutations using PolyPhen-2. Curr. Protoc. Hum. Genet. Chapter 7, Unit7.20 (2013).

85. Vaser, R., Adusumalli, S., Leng, S. N., Sikic, M. & Ng, P. C. SIFT missense predictions for genomes. Nat. Protoc. 11, 1–9 (2016).

86. Shihab, H. A. et al. Ranking non-synonymous single nucleotide polymorphisms based on disease concepts. Hum. Genomics 8, 11 (2014).

87. Lek, M. et al. Analysis of protein-coding genetic variation in 60,706 humans. Nature 536, 285–291 (2016).

88. Sherry, S. T. et al. dbSNP: the NCBI database of genetic variation. Nucleic Acids Res. 29, 308–311 (2001).

89. McLaren, W. et al. The Ensembl Variant Effect Predictor. Genome Biol. 17, 122 (2016).

90. Hubbard, T. et al. The Ensembl genome database project. Nucleic Acids Res. 30, 38–41 (2002).

91. Jeffares, D. C. et al. Transient structural variations have strong effects on quantitative traits and reproductive isolation in fission yeast. Nat. Commun. 8, 14061 (2017).

92. Chen, R., Im, H. & Snyder, M. Whole-Exome Enrichment with the Roche NimbleGen SeqCap EZ Exome Library SR Platform. Cold Spring Harb. Protoc. 2015, pdb.prot084855 (2015).

93. Li, H. & Durbin, R. Fast and accurate short read alignment with Burrows-Wheeler transform. Bioinforma. Oxf. Engl. 25, 1754–1760 (2009).

94. Talevich, E., Shain, A. H., Botton, T. & Bastian, B. C. CNVkit: Genome-Wide Copy Number Detection and Visualization from Targeted DNA Sequencing. PLoS Comput. Biol. 12, e1004873 (2016).

95. Belleau, P., Deschenes, A., Sun, G., Tuveson, D. A. & Krasnitz, A. CNprep:Copy Number Event Detection. (2020).

96. Li, H. & Durbin, R. Fast and accurate short read alignment with Burrows–Wheeler transform. Bioinformatics 25, 1754–1760 (2009).

97. Li, H. et al. The Sequence Alignment/Map format and SAMtools. Bioinforma. Oxf. Engl. 25, 2078–2079 (2009).

98. Barnett, D. W., Garrison, E. K., Quinlan, A. R., Strömberg, M. P. & Marth, G. T. BamTools: a C++ API and toolkit for analyzing and managing BAM files. Bioinformatics 27, 1691–1692 (2011).

99. Koboldt, D. C. et al. VarScan 2: somatic mutation and copy number alteration discovery in cancer by exome sequencing. Genome Res. 22, 568–576 (2012).

100. Wang, K., Li, M. & Hakonarson, H. ANNOVAR: functional annotation of genetic variants from high-throughput sequencing data. Nucleic Acids Res. 38, e164 (2010).

101. Landrum, M. J. et al. ClinVar: improving access to variant interpretations and supporting evidence. Nucleic Acids Res. 46, D1062–D1067 (2018).

102. Mayakonda, A., Lin, D.-C., Assenov, Y., Plass, C. & Koeffler, H. P. Maftools: efficient and comprehensive analysis of somatic variants in cancer. Genome Res. 28, 1747–1756 (2018).

103. Wang, Z. et al. SMASH, a fragmentation and sequencing method for genomic copy number analysis. Genome Res. 26, 844–851 (2016).

104. Gu, Z., Eils, R. & Schlesner, M. Complex heatmaps reveal patterns and correlations in multidimensional genomic data. Bioinforma. Oxf. Engl. 32, 2847–2849 (2016).

105. Belleau, P., Deschênes, A., Beyaz, S., Tuveson, D. A. & Krasnitz, A. CNVMetrics package: Quantifying similarity between copy number profiles.

106. Deschênes, A., Belleau, P., Tuveson, D. A. & Krasnitz, A. Quantifying similarity between copy number profiles with CNVMetrics package.

107. Zheng, G. X. Y. et al. Massively parallel digital transcriptional profiling of single cells. Nat. Commun. 8, 14049 (2017).

108. Frankish, A. et al. GENCODE reference annotation for the human and mouse genomes. Nucleic Acids Res. 47, D766–D773 (2019).

109. Edwards, N. et al. SCExecute: custom cell barcode-stratified analyses of scRNA-seq data. Bioinformatics 39, btac768 (2023).

110. Belleau, P., Deschênes, A., Chambwe, N., Tuveson, D. A. & Krasnitz, A. Genetic Ancestry Inference from Cancer-Derived Molecular Data across Genomic and Transcriptomic Platforms. Cancer Res. 83, 49–58 (2023).

111. Shen, R. & Seshan, V. E. FACETS: allele-specific copy number and clonal heterogeneity analysis tool for high-throughput DNA sequencing. Nucleic Acids Res. 44, e131 (2016).

112. Wang, W. et al. Single-cell transcriptomic atlas of the human endometrium during the menstrual cycle. Nat. Med. 26, 1644–1653 (2020).

113. Martinez-de-Morentin, X. et al. LIBRA: an adaptative integrative tool for paired single-cell multi-omics data. Quant. Biol. Beijing China 11, 246–259 (2023).

