## Supplemental files for "A High-Fidelity and Ancestrally Inclusive Patient-Derived Organoid Platform Resolves Cancer Cell Plasticity in Uterine Carcinosarcoma"

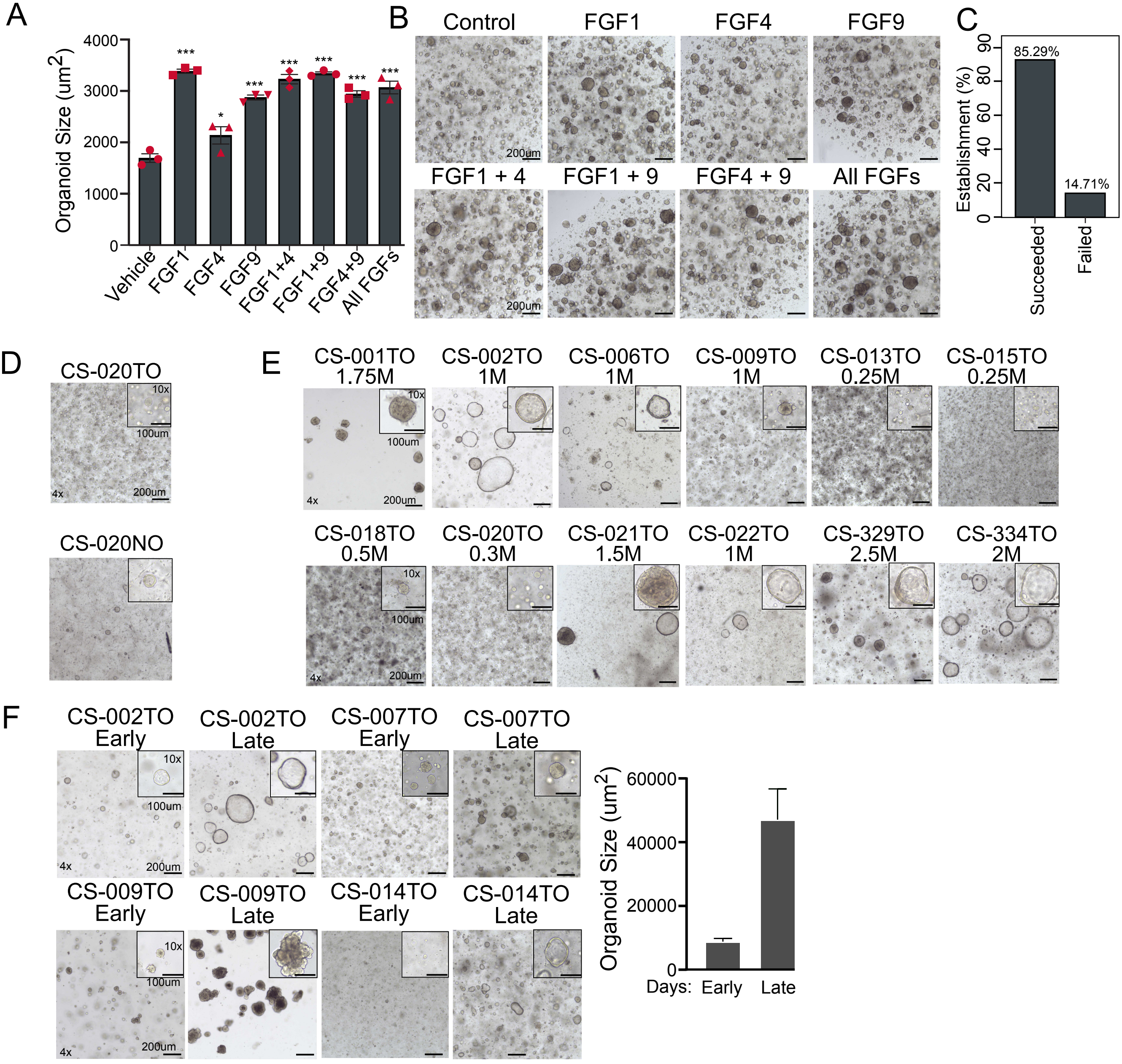
A
Organoid Size (um2)
4000
3000
2000
1000
0
***
*
Vehicle
FGF1
FGF4
FGF9
FGF1+4
FGF1+9
FGF4+9
All FGFs
B
Control
FGF1
FGF4
FGF9
FGF1 + 4
FGF1 + 9
FGF4 + 9
All FGFs
200um
C
Establishment (%)
85.29%
14.71%
Succeeded
Failed
D
CS-020TO
CS-020NO
10x
100um
4x
200um
E
CS-001TO
1.75M
CS-002TO
1M
CS-006TO
1M
CS-009TO
1M
CS-013TO
0.25M
CS-015TO
0.25M
CS-018TO
0.5M
CS-020TO
0.3M
CS-021TO
1.5M
CS-022TO
1M
CS-329TO
2.5M
CS-334TO
2M
F
CS-002TO
Early
CS-002TO
Late
CS-007TO
Early
CS-007TO
Late
CS-009TO
Early
CS-009TO
Late
CS-014TO
Early
CS-014TO
Late
Organoid Size (um2)
60000
40000
20000
0
Days:
Early
Late

**Figure S2:**

(**A**) Growth of UCS PDOs cultured with individual FGF ligands or combinations at indicated concentrations of 25 ng/mL (FGF1), 200 ng/mL (FGF9), 50 ng/mL (FGF4). Relative organoid formation efficiency was assessed at day 15 (mean ± SD, n = 3 (technical replicates), *p < 0.05, ***p < 0.001, Holm-Šídák's multiple comparisons test).

(**B**) Representative brightfield images of UCS PDOs cultured under indicated FGF conditions. Scale bar: x4: 200 µm.

(**C**) Establishment efficiency (%) of UCS PDOs is shown for each condition and defined as the proportion of successfully expanded lines.

(**D**) Representative brightfield images of additional matched normal endometrial PDOs and UCS PDOs from independent donors (n = 3). Scale bars: x4: 200 µm; x10: 100 µm.

(**E**) Long-term brightfield images of additional UCS PDO lines demonstrating sustained expansion over up to 2.5 months in FGF-enriched conditions. Scale bars, x4: 200 µm; x10: 100 µm.

(**F**) Representative brightfield images and quantification of PDO growth over time. PDO size ($\mu m^2$) was measured at indicated time points using ImageJ.

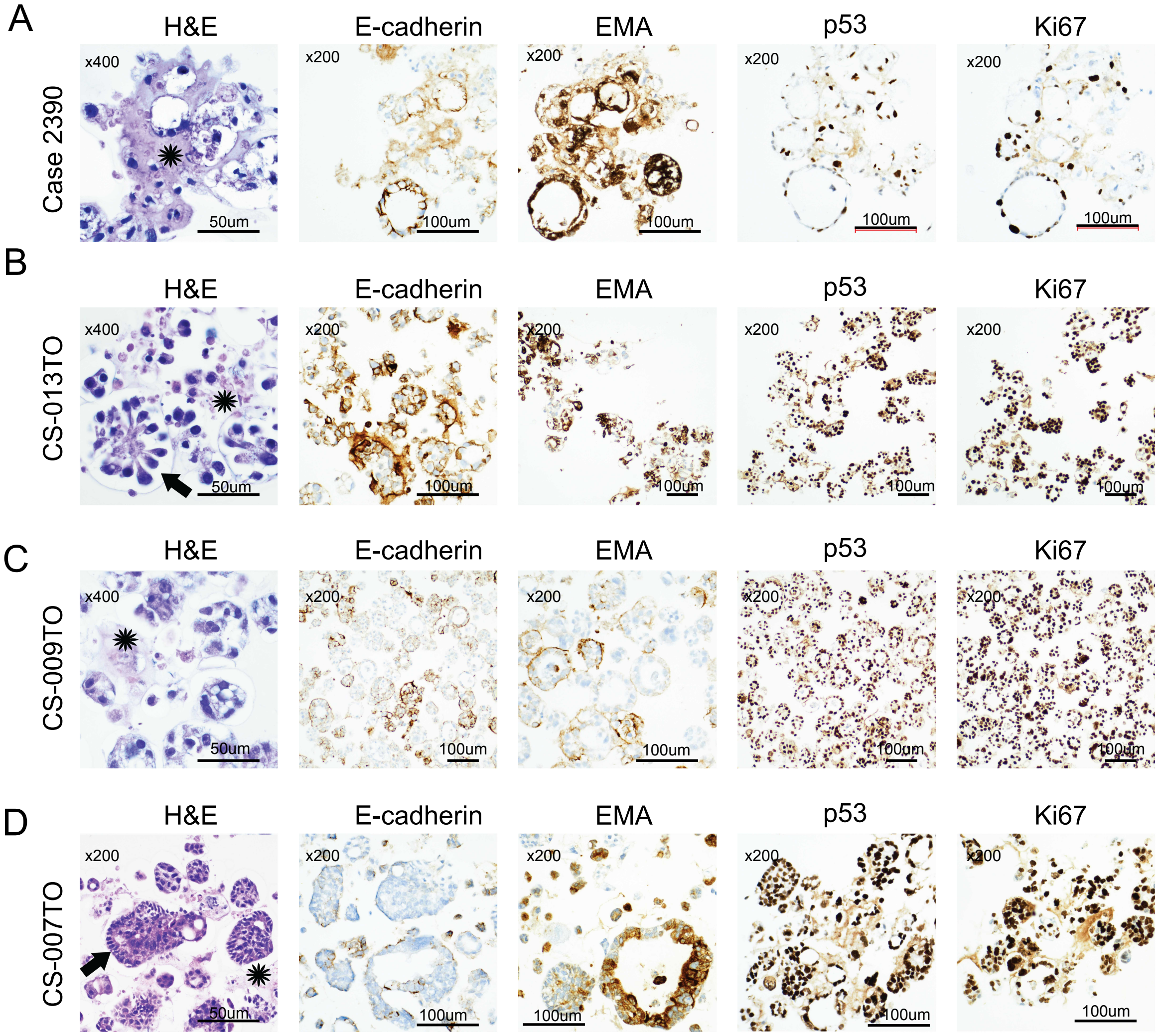

A
H&E
E-cadherin
EMA
p53
Ki67
Case 2390
x400
x200
50um
100um
B
CS-013TO
C
CS-009TO
D
CS-007TO

**Figure S3:**
(**A**) Additional representative H&E and IHC stainings (E-cadherin, EMA, p53, Ki67) of primary UCS tissue. Scale bars: 50 µm (H&E) and 100 µm (IHC). Area of interest marked *.
(**B**) Additional representative H&E and IHC stainings (E-cadherin, EMA, p53, Ki67) of USC PDO line CS-013TO. Scale bars: 50 µm (H&E) and 100 µm (IHC). Area of interest marked *. Arrows are indicating biphasic morphology.
(**C**) Additional representative H&E and IHC stainings (E-cadherin, EMA, p53, Ki67) of USC PDO line CS-009TO. Scale bars: 50 µm (H&E) and 100 µm (IHC). Area of interest marked *.
(**D**) Additional representative H&E and IHC stainings (E-cadherin, EMA, p53, Ki67) of USC PDO line CS-007TO. Scale bars: 50 µm (H&E) and 100 µm (IHC). Area of interest marked *. Arrows are indicating biphasic morphology.

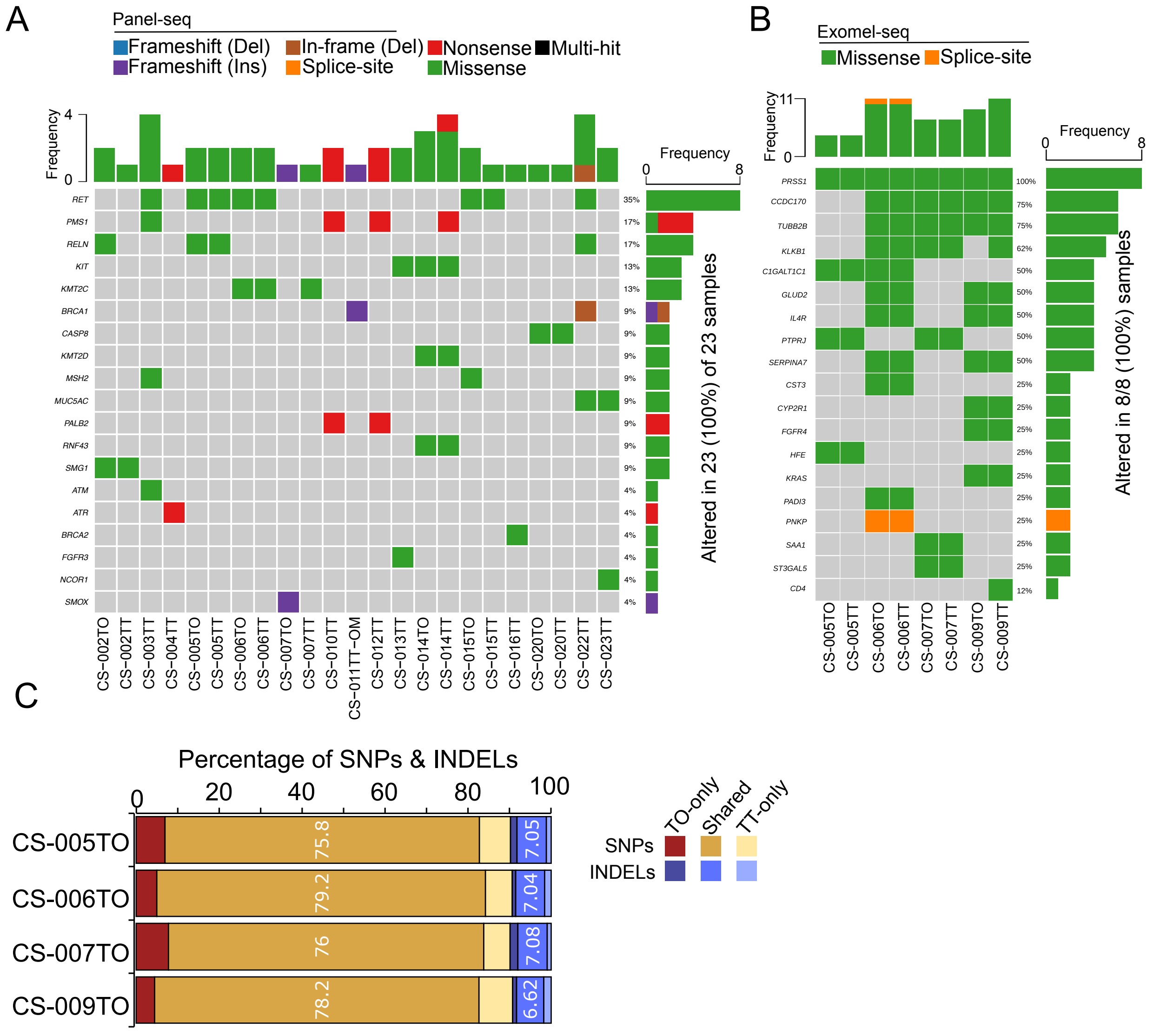

A
Panel-seq
Frameshift (Del)
In-frame (Del)
Nonsense
Multi-hit
Frameshift (Ins)
Splice-site
Missense
Frequency
RET
PMS1
RELN
KIT
KMT2C
BRCA1
CASP8
KMT2D
MSH2
MUC5AC
PALB2
RNF43
SMG1
ATM
ATR
BRCA2
FGFR3
NCOR1
SMOX
35%
17%
17%
13%
13%
9%
9%
9%
9%
9%
9%
9%
9%
4%
4%
4%
4%
4%
4%
Altered in 23 (100%) of 23 samples
CS−002TO CS−002TT CS−003TT CS−004TT CS−005TO CS−005TT CS−006TO CS−006TT CS−007TO CS−007TT CS−010TT CS−011TT−OM CS−012TT CS−013TT CS−014TO CS−014TT CS−015TO CS−015TT CS−016TT CS−020TO CS−020TT CS−022TT CS−023TT
B
Exomel-seq
Missense
Splice-site
Frequency
PRSS1
CCDC170
TUBB2B
KLKB1
C1GALT1C1
GLUD2
IL4R
PTPRJ
SERPINA7
CST3
CYP2R1
FGFR4
HFE
KRAS
PADI3
PNKP
SAA1
ST3GAL5
CD4
100%
75%
75%
62%
50%
50%
50%
50%
50%
25%
25%
25%
25%
25%
25%
25%
25%
25%
12%
Altered in 8/8 (100%) samples
CS-005TO CS-005TT CS-006TO CS-006TT CS-007TO CS-007TT CS-009TO CS-009TT
C
Percentage of SNPs & INDELs
0 20 40 60 80 100
CS-005TO
75.8
7.05
CS-006TO
79.2
7.04
CS-007TO
76
7.08
CS-009TO
78.2
6.62
TO-only
Shared
TT-only
SNPs
INDELs

**Figure S4:**

(**A**) Oncoplot showing targeted Panel-seq showing germline mutations across UCS tumors (n = 25; 23 primary, 2 metastases) and matched PDOs (n = 10 matched pairs). Mutation frequency per gene across matched pairs is indicated.

(**B**) Whole-exome sequencing (WES) derived oncoplot showing germline mutations across matched UCS primary tumor-organoid pairs (n = 4 donors; 8 samples total). Mutation frequency per gene across matched pairs is indicated.

(**C**) Percentage distribution of shared and sample-specific germline variants between primary tumors (TT) and matched UCS PDOs (TO). Stacked bar plots show proportions of shared germline SNPs and insertions/deletions (INDELs), as well as tumor-only and organoid-only variants for each donor. Across donors, most variants were shared between primary tumors and PDOs (75.8–79.2%). Variants were classified as shared if detected in both tumor and PDOs.

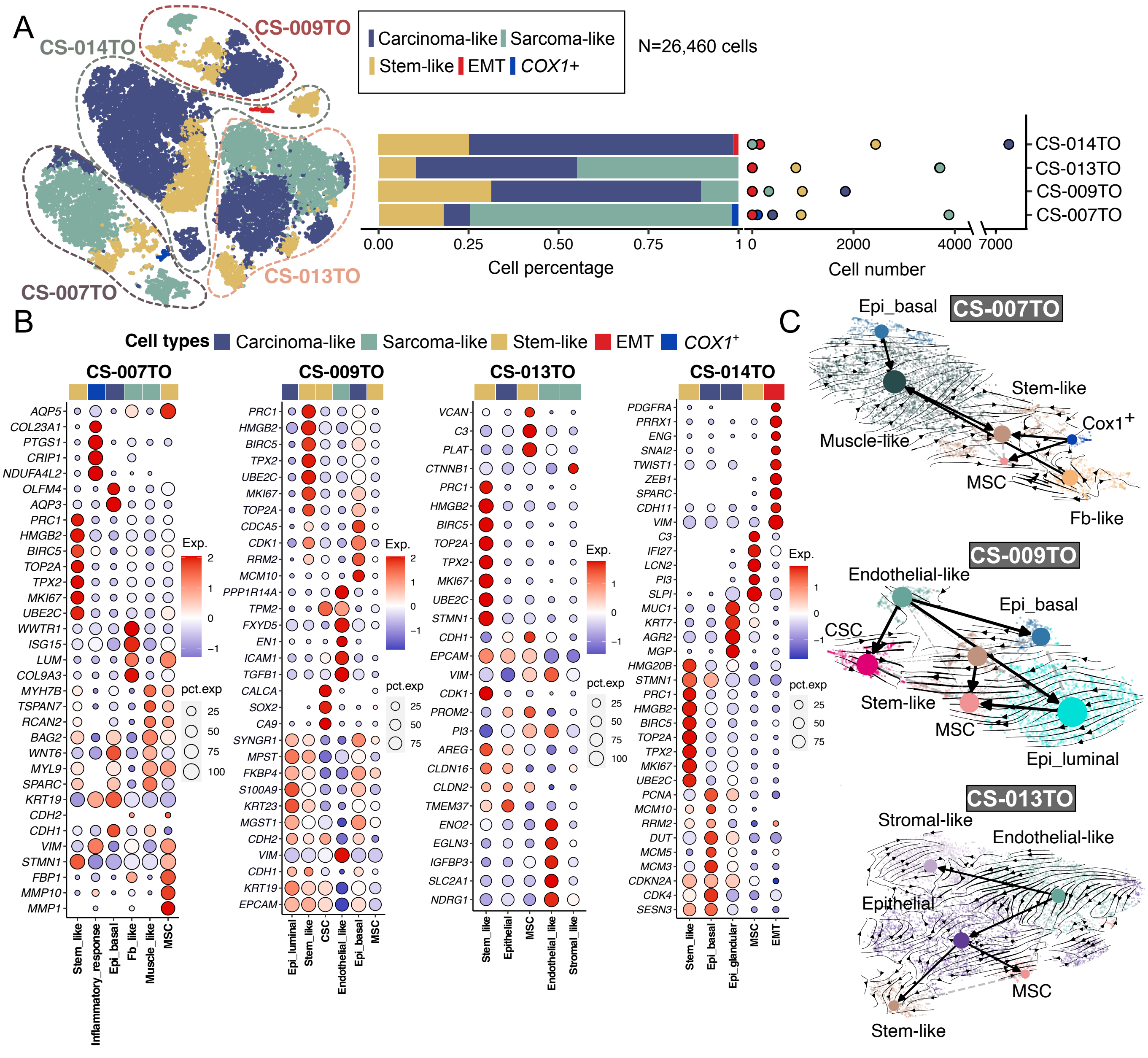
A
CS-014TO
CS-009TO
CS-007TO
CS-013TO
Carcinoma-like Sarcoma-like
Stem-like EMT COX1+
N=26,460 cells
CS-014TO
CS-013TO
CS-009TO
CS-007TO
0.00 0.25 0.50 0.75 1
Cell percentage
0 2000 4000 7000
Cell number
B
Cell types Carcinoma-like Sarcoma-like Stem-like EMT COX1+
CS-007TO
AQP5 COL23A1 PTGS1 CRIP1 NDUFA4L2 OLFM4 AQP3 PRC1 HMGB2 BIRC5 TOP2A TPX2 MKI67 UBE2C WWTR1 ISG15 LUM COL9A3 MYH7B TSPAN7 RCAN2 BAG2 WNT6 MYL9 SPARC KRT19 CDH2 CDH1 VIM STMN1 FBP1 MMP10 MMP1
Stem_like Inflammatory_response Epi_basal Fb_like Muscle_like MSC
Exp. 2 1 0 −1
pct.exp 25 50 75 100
CS-009TO
PRC1 HMGB2 BIRC5 TPX2 UBE2C MKI67 TOP2A CDCA5 CDK1 RRM2 MCM10 PPP1R14A TPM2 FXYD5 EN1 ICAM1 TGFB1 CALCA SOX2 CA9 SYNGR1 MPST FKBP4 S100A9 KRT23 MGST1 CDH2 VIM CDH1 KRT19 EPCAM
Epi_luminal Stem_like CSC Endothelial_like Epi_basal MSC
Exp. 2 1 0 −1
pct.exp 25 50 75
CS-013TO
VCAN C3 PLAT CTNNB1 PRC1 HMGB2 BIRC5 TOP2A TPX2 MKI67 UBE2C STMN1 CDH1 EPCAM VIM CDK1 PROM2 PI3 AREG CLDN16 CLDN2 TMEM37 ENO2 EGLN3 IGFBP3 SLC2A1 NDRG1
Stem_like Epithelial MSC Endothelial_like Stromal_like
Exp. 1 0 −1
pct.exp 25 50 75 100
CS-014TO
PDGFRA PRRX1 ENG SNAI2 TWIST1 ZEB1 SPARC CDH11 VIM C3 IFI27 LCN2 PI3 SLPI MUC1 KRT7 AGR2 MGP HMG20B STMN1 PRC1 HMGB2 BIRC5 TOP2A TPX2 MKI67 UBE2C PCNA MCM10 RRM2 DUT MCM5 MCM3 CDKN2A CDK4 SESN3
Stem_like Epi_basal Epi_glandular MSC EMT
Exp. 1 0 −1
pct.exp 25 50 75
C
CS-007TO
Epi_basal
Stem-like
Cox1+
Muscle-like
MSC
Fb-like
CS-009TO
Endothelial-like
Epi_basal
CSC
Stem-like
MSC
Epi_luminal
CS-013TO
Stromal-like
Endothelial-like
Epithelial
MSC
Stem-like

**Figure S5:**

(**A**) t-SNE representation of 51,784 total cells from scRNA-seq data of UCS PDOs (n = 4), grouped into major compartments: carcinoma-like, sarcoma-like, stem-like, and EMT-/*COX1*$^{+}$-associated populations. Stacked bar plot (right) shows proportional representation of each cell type per donor and respective cell numbers.

(**B**) Dot plots showing expression of representative marker genes used for annotation of cell types across subclusters of cell compartments for each donor. Dot size represents the percentage of cells expressing the gene within a cluster, and color intensity represents scaled average expression (equivalent to Z-score). Gene markers were predicted using FindAllMarkers from Seurat based on with adjusted $p < 0.05$ from Wilcoxon rank-sum test and log2 fold difference of 1.

(**C**) RNA velocity-inferred trajectories for additional PDO donors (CS-007TO, CS-009TO, CS-013TO). Directional flow suggests dynamic transitions between epithelial, stem-like, and mesenchymal populations across independent organoid lines. Velocity vectors were computed using scVelo based on spliced and unspliced transcript counts.

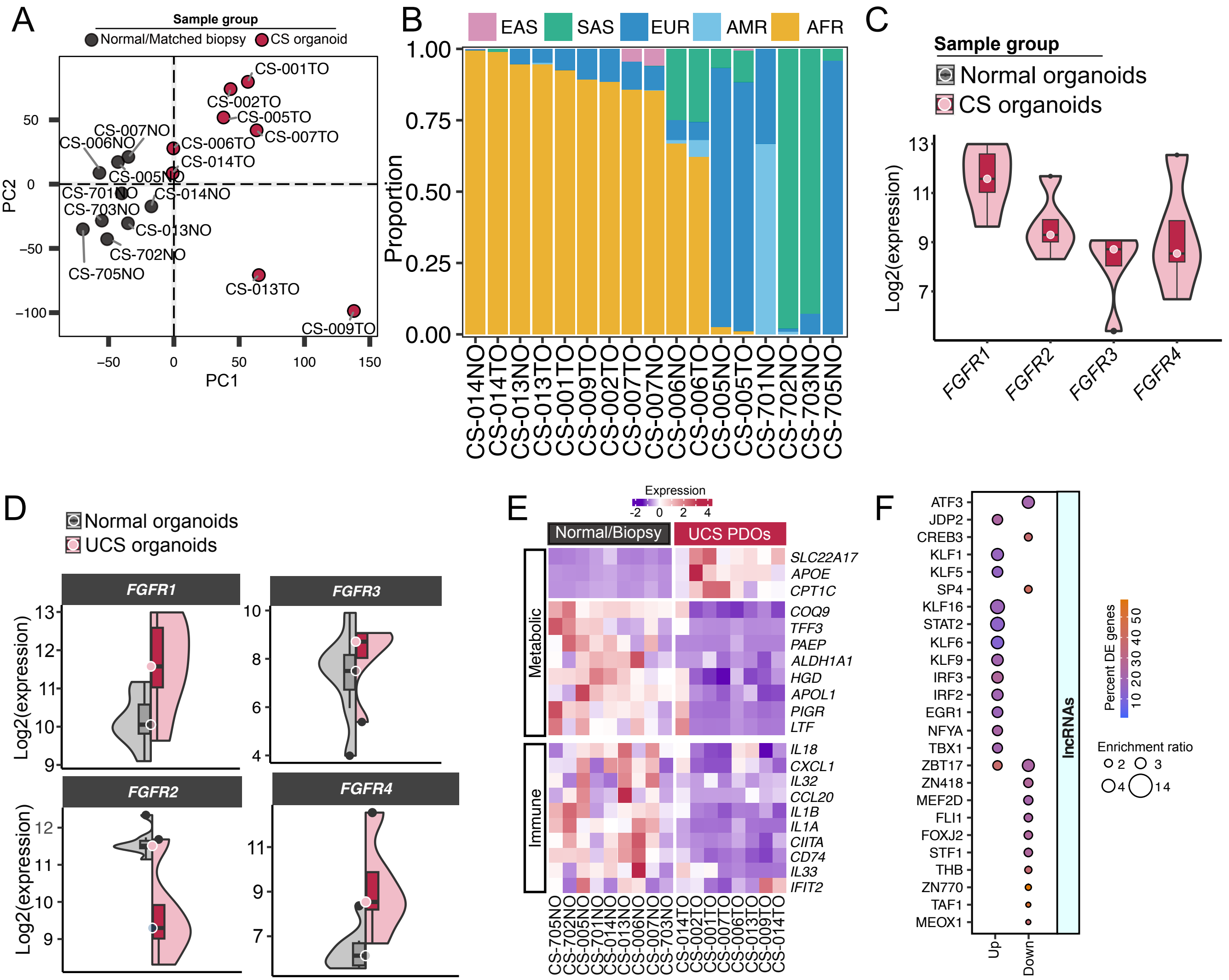

A
Sample group
Normal/Matched biopsy
CS organoid
CS-001TO
CS-002TO
CS-005TO
CS-007TO
CS-006TO
CS-014TO
CS-007NO
CS-006NO
CS-005NO
CS-701NO
CS-014NO
CS-703NO
CS-013NO
CS-702NO
CS-705NO
CS-013TO
CS-009TO
PC1
PC2
50
0
−50
−100
−50
0
50
100
150
B
EAS
SAS
EUR
AMR
AFR
Proportion
1.00
0.75
0.50
0.25
0.00
CS-014NO
CS-014TO
CS-013NO
CS-013TO
CS-001TO
CS-009TO
CS-002TO
CS-007TO
CS-007NO
CS-006NO
CS-006TO
CS-005NO
CS-005TO
CS-701NO
CS-702NO
CS-703NO
CS-705NO
C
Sample group
Normal organoids
CS organoids
Log2(expression)
13
11
9
7
FGFR1
FGFR2
FGFR3
FGFR4
D
Normal organoids
UCS organoids
FGFR1
FGFR3
FGFR2
FGFR4
Log2(expression)
13
12
11
10
9
10
8
6
4
12
11
10
9
11
9
7
E
Expression
-2
0
2
4
Normal/Biopsy
UCS PDOs
Metabolic
Immune
SLC22A17
APOE
CPT1C
COQ9
TFF3
PAEP
ALDH1A1
HGD
APOL1
PIGR
LTF
IL18
CXCL1
IL32
CCL20
IL1B
IL1A
CIITA
CD74
IL33
IFIT2
CS-705NO
CS-702NO
CS-005NO
CS-701NO
CS-014NO
CS-013NO
CS-006NO
CS-007NO
CS-703NO
CS-014TO
CS-002TO
CS-001TO
CS-007TO
CS-006TO
CS-013TO
CS-009TO
CS-014TO
F
ATF3
JDP2
CREB3
KLF1
KLF5
SP4
KLF16
STAT2
KLF6
KLF9
IRF3
IRF2
EGR1
NFYA
TBX1
ZBT17
ZN418
MEF2D
FLI1
FOXJ2
STF1
THB
ZN770
TAF1
MEOX1
Up
Down
lncRNAs
Percent DE genes
10
20
30
40
50
Enrichment ratio
2
3
4
14

**Figure S6:**

(**A**) Principal component analysis (PCA) of bulk RNA-seq data from UCS (n = 8) and normal endometrial biopsy-derived PDOs (n = 9). PCA was performed using CPM (counts per million) normalized counts from DESeq2.

(**B**) Ancestry inferred from bulk RNA-seq generated from UCS tumors and biopsy samples using ADMIXTURE analysis. Bar plots show estimated ancestry proportions (EAS, SAS, EUR, AMR, AFR) per donor.

(**C**) Violin plots showing expression levels (log2 CPM normalized counts) of FGFR family members (FGFR1-4) in UCS.

(**D**) Violin plots showing differential expression levels (log2 CPM normalized counts) of FGFR family members (FGFR1-4) in UCS compared to normal PDOs. Indicated genes were significantly differentially expressed with log2 fold difference ± 1 and adjusted p-value < 0.05 derived from DESeq2 (except non-significant *FGFR3*).

(**E**) Heatmap of showing expression patterns of the genes associated with metabolic and immune-related pathways between UCS and normal PDOs. Gene expression values are Z-score-scaled log2 CPM normalized counts.

(**F**) Promoter motif enrichment analysis of upregulated and downregulated long non-coding RNAs (lncRNAs) in UCS PDOs relative to NO controls. Motif enrichment was assessed using SEA (Simple Enrichment Analysis) from MEME Suite, revealing significant enrichment of CREB-family transcription factor binding motif (CREB3) in the downregulated gene promoters. Enrichment ratio was calculated using ratio of true positives and false positive sequences determined by SEA. The color gradient represents percentage of differentially expressed genes in UCS regulated by corresponding transcription factors.

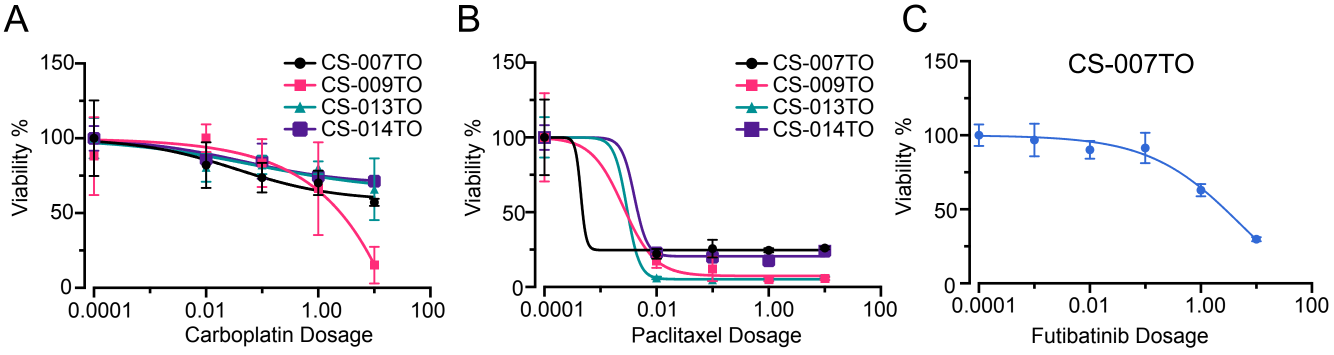

A
B
C
150
100
50
0
Viability %
CS-007TO
CS-009TO
CS-013TO
CS-014TO
0.0001
0.01
1.00
100
Carboplatin Dosage
Paclitaxel Dosage
CS-007TO
Futibatinib Dosage

**Figure S7:**

(**A, B**) Dose-response curves of UCS PDO (n = 4) treated with Carboplatin (**A**) and Paclitaxel (**B**) (0.01-10 µM). PDOs were treated for 6 days, and viability was quantified using the CellTiter-Glo 3D luminescent assay. Data is normalized to vehicle control and presented as mean ± SD from at least 3 technical replicates.

(**C**) Dose-response curves of UCS PDO CS-007TO treated with Futibatinib (0.001-10 µM). PDOs were treated for 6 days, and viability was quantified using the CellTiter-Glo 3D luminescent assay (n > 4 technical replicates).
